# The evolutionary conserved choroid plexus sustains the homeostasis of brain ventricles in zebrafish

**DOI:** 10.1101/2023.11.03.565468

**Authors:** Inyoung Jeong, Søren N. Andreassen, Linh Hoang, Morgane Poulain, Yongbo Seo, Hae-Chul Park, Maximilian Fürthauer, Nanna MacAulay, Nathalie Jurisch-Yaksi

## Abstract

The choroid plexus produces cerebrospinal fluid (CSF) by transport of electrolytes and water from the vasculature to the brain ventricles. The choroid plexus plays additional roles in brain development and homeostasis by secreting neurotrophic molecules, and by serving as a CSF-blood barrier and immune interface. Prior studies have identified transporters on the epithelial cells that transport water and ions into the ventricles and tight junctions involved in the CSF-blood barrier. Yet, how the choroid plexus epithelial cells maintain the brain ventricle system and control brain physiology remain unresolved. To provide novel insights into the physiological roles of the choroid plexus, we use juvenile and adult zebrafish as model systems. Upon histological and transcriptomic analyses, we first identified that the zebrafish choroid plexus is highly conserved with the mammalian choroid plexus and that it expresses all transporters necessary for CSF secretion. Using novel genetic lines, we also identified that the choroid plexus secretes proteins into the CSF. Next, we generated a transgenic line allowing us to ablate specifically the epithelial cells in the choroid plexus. Using the ablation system, we identified a reduction of the ventricular sizes, but no alterations of the CSF-blood barrier. Altogether, our findings identified that the zebrafish choroid plexus is evolutionarily conserved and critical for maintaining the size and homeostasis of the brain ventricles.

**Graphical abstract:** 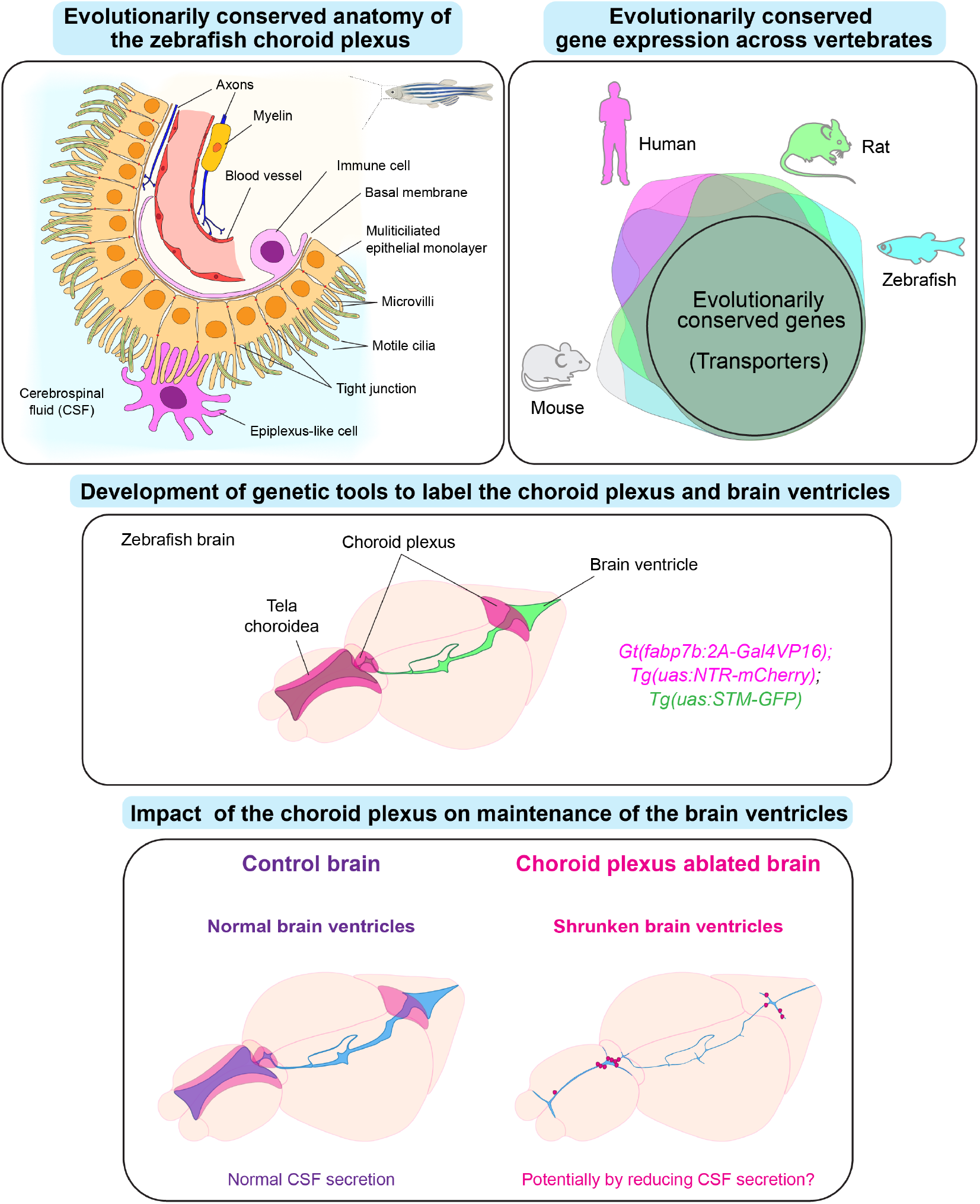

**Highlights:**

- The zebrafish choroid plexus has similar anatomical features with the mammalian choroid plexus.
- The expression of choroid plexus transporters involved in CSF secretion is evolutionarily conserved across vertebrates.
- Generation of a novel choroid plexus specific driver line shows that the choroid plexus epithelial cells secrete proteins into CSF.
- Ablation of the choroid plexus decreases the size of the brain ventricles.

## Introduction

The choroid plexus (ChP), which produces cerebrospinal fluid (CSF), is a secretory epithelium located in the brain ventricles (MacAulay *et al*, 2022). The ChP consists of multiple cell types including mainly ciliated epithelial cells, but also endothelium, immune cells, glia and neurons (Dani *et al*, 2021). Epithelial cells of the ChP generate CSF by transepithelial transport of electrolytes and fluid from the vasculature to the ventricular compartment (MacAulay *et al*., 2022). The ChP also secretes neurotrophic and growth factors into CSF that support embryonic and postnatal neurodevelopment, and brain homeostasis (Lun *et al*, 2015b; Saunders *et al*, 2023). Thanks to tight junctions located between epithelial cells, the ChP forms the CSF-blood barrier, which is a barely permeable border between CSF and blood vessels necessary to maintain brain homeostasis (Solar *et al*, 2020). Finally, the ChP plays an important role for immune surveillance of the brain (Cui *et al*, 2020; Kunis *et al*, 2013; Shipley *et al*, 2020). Due to its multiple roles in the brain, alterations of the ChP are associated with various neurological diseases. For instance, disturbed water homeostasis such as CSF hypersecretion by the ChP may contribute to posthemorrhagic hydrocephalus (Ben-Shoshan *et al*, 2023; Hochstetler *et al*, 2022; Karimy *et al*, 2017; Lolansen *et al*, 2022). ChP abnormalities related to barrier dysfunction, immune response and CSF flow have also been linked with neurodegenerative diseases (Kaur *et al*, 2016; Liu *et al*, 2022; Solar *et al*., 2020).

Although numerous studies have investigated the molecular mechanisms underlying CSF secretion by the ChP in physiological and pathological conditions (Bothwell *et al*, 2019; Kaur *et al*., 2016; MacAulay *et al*., 2022; Saunders *et al*., 2023), how the choroid plexus maintains the brain ventricular system and thereby regulates brain physiology remain poorly understood. We trust that the zebrafish offers opportunities to address this knowledge gap due to its smaller yet conserved brain, well characterized neural circuits, and amenable genetic tools (Jurisch-Yaksi *et al*, 2020; Wyatt *et al*, 2015). To date, however, studies in zebrafish have mainly focused on the development of the ChP. Prior work has revealed that zebrafish have two ChPs, which are located on the dorsal roof of the forebrain and hindbrain (Garcia-Lecea *et al*, 2008; Korzh, 2023). The ChPs appear during early embryonic stage (at circa 36 hour-post-fertilization (hpf)) (Garcia-Lecea *et al*., 2008) and are composed of motile monociliated epithelial cells before eventually becoming a multiciliated organ at juvenile stage (D’Gama *et al*, 2021; Olstad *et al*, 2019). No study has however reported the physiological properties of ChP in the zebrafish, limiting our understanding of its contribution to CSF production, immune surveillance, CSF-blood barrier, and brain homeostasis.

To fill this knowledge gap, we performed histological, transcriptomic, and physiological analyses of the ChP. In our study, we first report evolutionary conserved anatomical and molecular features of the zebrafish ChP, compared to mammals. We then developed novel genetic tools to label and manipulate the ChP and visualize the brain ventricles. Using these tools, we genetically ablated ChP epithelial cells and identified that the ChP plays a crucial for CSF secretion and maintenance of the brain ventricles. By providing an extensive analysis of ChP in the zebrafish, we trust that our work set the basis for a future understanding of ChP function in health and disease of the brain.

## Results

### The zebrafish choroid plexus shows evolutionary conserved anatomical characteristics

To characterize the tissue architecture of the zebrafish choroid plexus, we performed histological analyses of both the forebrain ChP (Fig. 1A-B) and hindbrain ChP (Fig. 1C-D) in wholemount adult brain explants. We focused on key characteristics of the ChP that were previously described in mammals, namely the organization of epithelial cells, their proximity to CSF and blood vessels, the presence of immune cells and the innervation by neurons.

**Figure 1.**
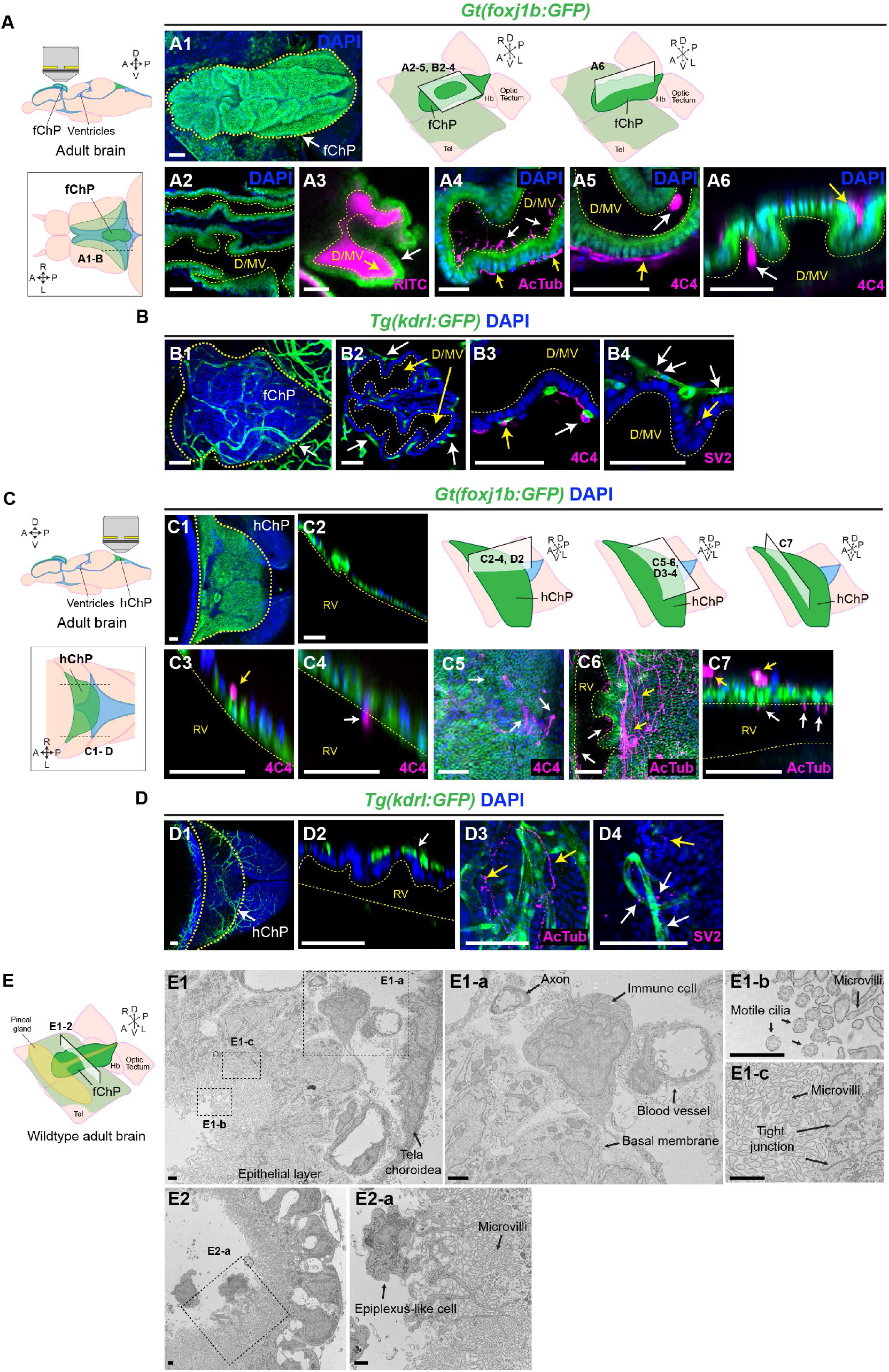
The zebrafish choroid plexuses show evolutionarily conserved tissue characteristics. **(A)** Diagram of the adult brain explant for imaging the forebrain choroid plexus (fChP) and di-/mesencephalic ventricle (D/MV). Lateral (upper) and dorsal (lower) views of the brain. **(A1-A6)** *Gt(foxj1b:GFP)* labels ChP epithelial cells. DAPI labels nuclei. Yellow dotted lines mark either fChP or D/MV. **(A1)** Max projection of the fChP (left) and schemes representing the location of the images shown in (A2-B4) (right). **(A2)** Single plane of the fChP showing the folded epithelium. **(A3)** Single plane of the fChP injected with 70kDa RITC-dextran (magenta) into the telencephalic ventricle. **(A4)** Single plane of the fChP immunolabelled by acetylated tubulin (AcTub) antibody marking both cilia (white arrows) and axons (yellow arrows). **(A5-A6)** Single plane of the fChP labelled with 4C4 antibody marking immune cells. **(B1-B4)** Max projection **(B1)** and single plane **(B2-B4)** of the fChP in the *Tg(kdrl:GFP)* that labels blood endothelial cells (white arrows) immunolabelled by GFP, 4C4 for immune cells **(B3)** and SV2 antibodies for presynaptic vesicles **(B4)**. DAPI labels nuclei. **(C)** Diagram of the adult brain explant for imaging the hindbrain choroid plexus (hChP) and rhombencephalic ventricle (RV). Lateral (upper) and dorsal (lower) views of the brain. **(C1-C7)** *Gt(foxj1b:GFP)* labels ChP epithelial cells. DAPI labels nuclei. Yellow dotted lines mark either hChP or RV. **(C1)** Max projection of the hChP. **(C2)** Single plane of the hChP showing the sheet-like epithelium (left) and schemes representing the location of the images shown in C2-D4 (right). **(C3-C5)** Single plane **(C3-C4)** and max projection **(C5)** images of the hChP labelled with 4C4 antibody. **(C6-C7)** Max projection **(C6)** and single plane **(C7)** of the hChP labelled with AcTub antibody. **(D1-D4)** Max projection **(D1, D3-D4)** and single plane **(D2)** images of the hChP in the *Tg(kdrl:GFP)* immunolabelled by GFP **(D2)**, AcTub **(D3)** and SV2 antibodies **(D4)**. DAPI labels nuclei. **(E)** Diagram of the fChP in the adult brain explant used for transmission electron microscopy. **(E1, E2)** Cross section view images of the fChP by transmission electron microscope. **(E1-a-E1-c, E2-a)** High magnification images from dotted boxes from (E1) or (E2). Abbreviations: A, anterior; P, posterior; D, dorsal; V, ventral; R, right; L, left; Hb, habenula; Tel, telencephalon. Scale bars: (A1-D4), 50 µm; (E1-E2a): 1 µm.

To specifically label epithelial cells of the ChP, we used the *Gt(foxj1b:GFP)* line (Tian *et al*, 2009), which expressed GFP in the forebrain ChP (D’Gama *et al*., 2021; Olstad *et al*., 2019). Using this line, we observed that both the forebrain and hindbrain ChP are composed mainly of *foxj1b*-expresssing ciliated epithelial cells which are organized in a polarized monolayer (Fig. 1). Interestingly, while the forebrain ChP is highly folded around multiple connected cavities filled with CSF (Fig. 1A1-3), the hindbrain ChP is arranged in a sheet of cells located on top of the ventricle (Fig. 1C1-2).

**Figure 2.**
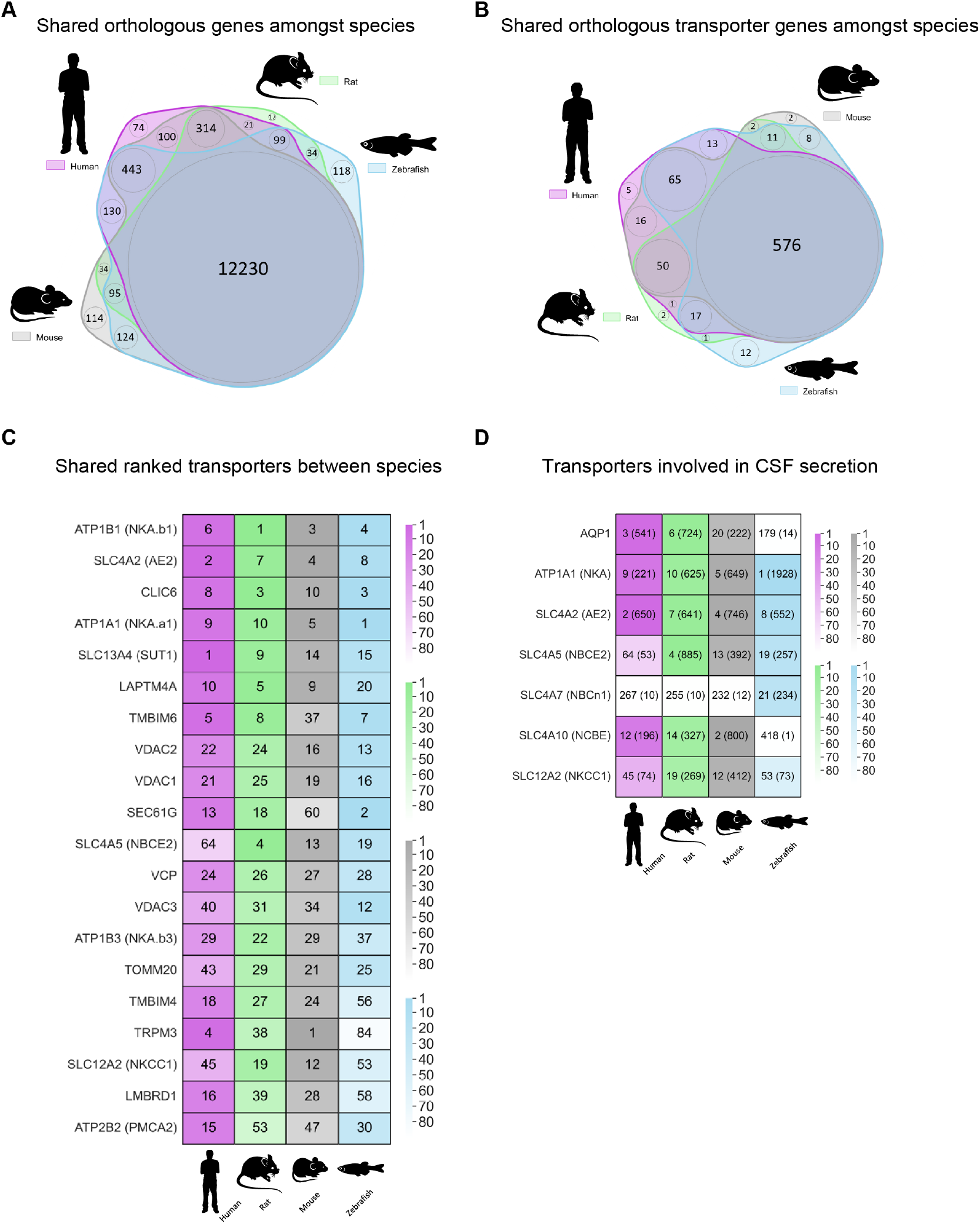
Expression of transporter genes is evolutionarily conserved across vertebrates. **(A)** Weighted Venn diagram showing protein coding genes expressed in the ChP that have an ortholog in human, rat, mouse and zebrafish. **(B)** Weighted Venn diagram showing transporter genes expressed in the ChP that have an ortholog in all four species. **(C)** Table of the ranked expression of transporter genes for each species, sorted based on the mean rank of all four species. Darker colors indicate higher ranks. **(D)** Table of the ranked expression of transporter genes involved in cerebrospinal fluid (CSF) secretion. Darker colors indicate higher ranks. The numbers in parentheses indicate transcript per million (TPM).

**Figure 3.**
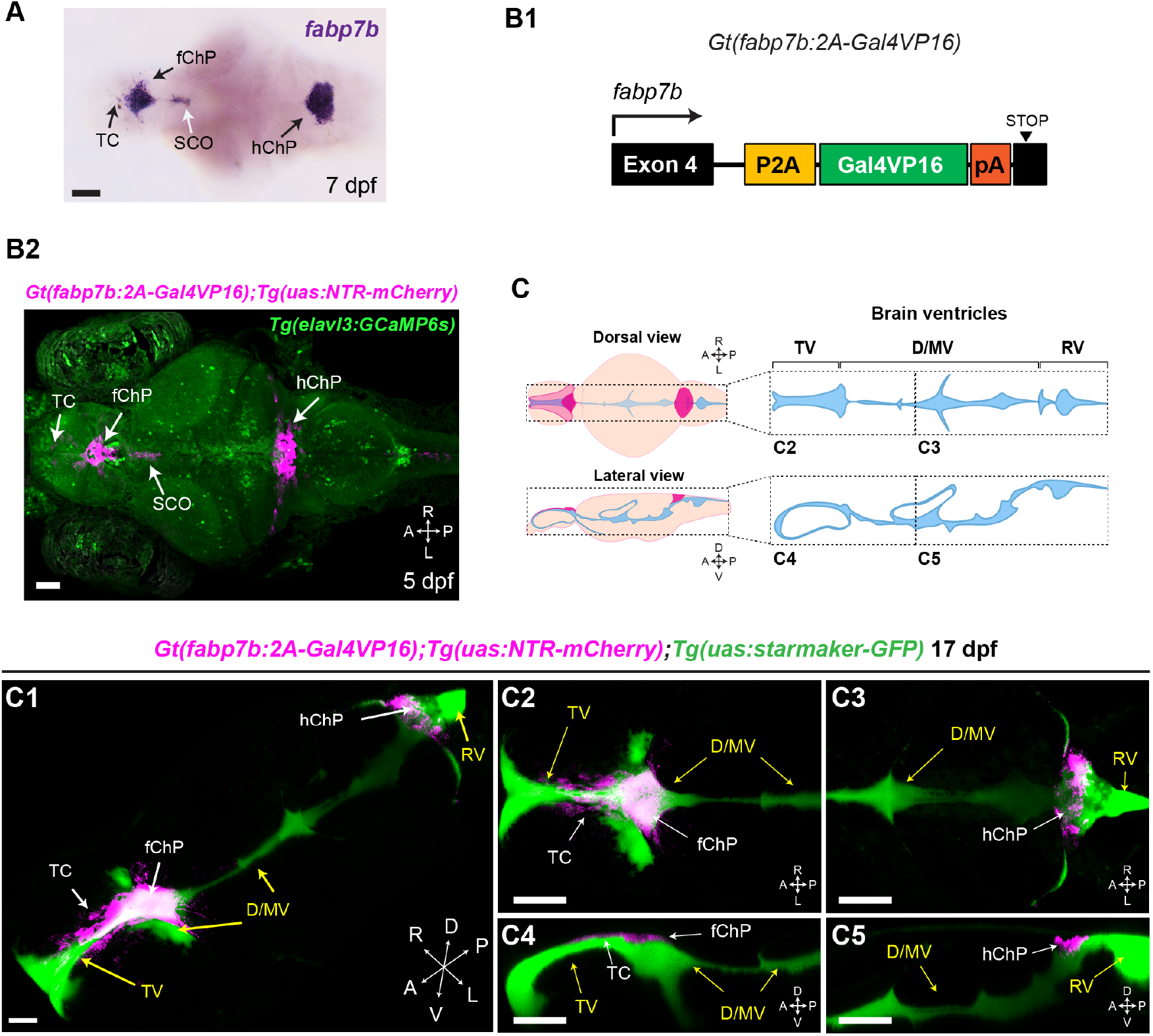
The fabp7b-expressing choroid plexus cells secrete proteins into the CSF and brain ventricles. **(A)** *fabp7b RNA* expression in the larval brain at 7 dpf (dorsal view) **(B1)** Schematic of the Gal4 integration in the *Gt(fabp7b:2A-Gal4vp16)* line. **(B2)** Dorsal view of the *Gt(fabp7b:2A-Gal4vp16);Tg(uas:NTR-mCherry);Tg(elavl3:GCaMP6s)* line at 5 dpf. **(C)** Diagrams of the brain parenchyma and ventricles of a juvenile (17-18dpf). The dotted boxes indicate the location of the images of brain ventricles in (C2-C5). **(C1-C5)** Images of *Gt(fabp7b:2A-Gal4vp16);Tg(uas:NTR-mCherry);Tg(uas:starmaker-GFP)* line at 17 dpf. **(C1)** 3D image reconstructed from confocal stacks. **(C2-C5)** Max projection images showing dorsal and sagittal view of the anterior and posterior part of the brain as indicated in (C). Abbreviations: TC, tela choroidea; fChP, forebrain choroid plexus; hChP, hindbrain choroid plexus; SCO, subcommisural organ; TV, telencephalic ventricle; D/MV, dien-/mesencephalic ventricle; RV: rhombencephalic ventricle. A, anterior; P, posterior; D, dorsal; V, ventral; R, right; L, left. Scale bars: 50 µm

Next, we examined whether the zebrafish ChP is vascularized to the same extent as the mammalian ChP. Using an endothelial specific transgenic line *Tg(kdrl:GFP)* (Beis et al, 2005), we observed many blood vessels near the epithelial cell layer of the forebrain (Fig.1B) and hindbrain ChP (Fig.1D). Importantly, blood vessels were found exclusively in the basolateral side of epithelial cells in both ChP (Fig.1B2-B4, 1D2), similarly to mammals.

Since the ChP plays important roles for immune surveillance (Cui *et al*., 2020; Kunis *et al*., 2013; Shipley *et al*., 2020), we investigated whether immune cells were present in the ChP. To mark immune cells, we performed immunostaining with the 4C4 antibody detecting galectin 3 binding protein b (lgals3bpb) (Rovira *et al*, 2023). We found several immune cells at the basolateral side of the forebrain (Fig.1A5, A6, B3) and hindbrain ChP (Fig.1C3-C4). Immune cells had long processes contacting multiple epithelial cells (Fig.1A5, yellow arrow, Fig.1C5). Immune cells were also adjacent to the blood vessels (Fig.1B3) but were contacting blood vessels to a lesser extent than epithelial cells. We also identified 4C4-positive cells at the apical surface of the forebrain and hindbrain ChP in the CSF-filled cavities (Fig.1A5, A6, C4), which may correspond to epiplexus cells, a specialized type of immune cells in the mammalian ChP (Carpenter *et al*, 1970; Peters, 1974).

Nerve innervations have been described in the mammalian ChP (Edvinsson *et al*, 1987; Edvinsson *et al*, 1974; Lindvall *et al*, 1978). To identify whether innervation is a conserved feature of the ChP, we labelled axons with an acetylated tubulin antibody. Note that acetyl-tubulin also labels cilia of epithelial cells (Fig.1A4 and C7, white arrows). We found axonal projections in the basolateral part of epithelial cells in both ChPs (Fig.1A4 and C7, yellow arrow). Next, by staining synaptic vesicle protein 2 (SV2), we noticed that pre-synaptic vesicles were more often in proximity to blood vessels (Fig.1B4 and D4, white arrows) than epithelial cells (Fig.1B4, yellow arrow).

To further investigate the ultrastructure of the ChP, we performed electron microscopy of the forebrain ChP (Fig.1E and Supple Fig.1). Like in our immunohistochemical analyses, we observed a monolayer of epithelial cells, blood vessels and axonal projections (Fig.1E1, 1E1-a). The epithelial layer had a clear polarized structure (Fig.1E1). In the apical part, which is in direct contact with CSF, we identified cilia with a 9+2 configuration and a large number of microvilli (Fig.1E1, 1E1-b), supporting a secretory role of the ChP. We also observed tight junctions between epithelial cells (Fig.1E1-c), which are required for the CSF-blood barrier (Vorbrodt & Dobrogowska, 2003). In the basolateral part, we noticed a basal membrane layer which contacted the blood vessels, immune cells and axons (Fig.1E1, 1E1-a). We also detected an epiplexus-like cell within the CSF filled cavity which interacted closely with microvilli on the epithelial cells (Fig.1E2, 1E2-a).

Next, we determined the anatomy of the midline tela choroidea, which we previously showed to be enriched in motile ciliated ependymal cells(D’Gama *et al*., 2021). We were especially interested to identify whether it has similar characteristics to the ChP (SuppleFig. 2) since there is no ChP described in the telencephalic ventricle of the zebrafish. We observed that, like the ChP, the tela choroidea is a multiciliated epithelial monolayer (SuppleFig. 2A3, 2A5, 2B1) with numerous microvilli, tight junctions (SuppleFig.2C2), vasculature, immune cells, axonal projections and synaptic vesicles (SuppleFig.2A2-D2). These data show that the tela choroidea has comparable tissue characteristics as the ChP and suggest that the tela choroidea may play a similar role as the ChP in the telencephalic ventricle.

Altogether, our analyses identified that the zebrafish ChP has evolutionarily conserved anatomical features.

### The choroid plexus develops throughout juvenile stages

We previously reported that the choroid plexus undergoes a transition from a monociliated to a multiciliated epithelium at juvenile stage (circa 3 weeks) when the brain ventricles expand significantly (D’Gama *et al*., 2021). This motivated us to identify whether the ChP features are already established at 15-17 days post fertilization, at the onset of multiciliation (SuppleFig.3B1-a). Notably at this stage, the forebrain ChP is organized in a flat epithelium which does not have folded cavities (SuppleFig.3A1-A3), and thereby resemble more the hindbrain ChP (SuppleFig.3C1-C2).

To visualize blood vessels, we microinjected Alexa-647 10 kDa dextran fluorescent dye into the cardinal vein of *Gt(foxj1b:GFP)* (SuppleFig.3A1, 3C1). Like in adults, we observed that blood vessels were located in the basolateral side of epithelial cells in the tela choroidea-ChPs (SuppleFig.3A1-a, -b, 3C1-a, -b, blue arrows). We also identified fluorescent signals in the brain ventricles (SuppleFig.3A1-b, 3C1-b, yellow dotted lines) upon microinjection of a 10 kDa dye in the cardinal vein.

Next, we detected 4C4-positive immune cells in both the apical and basolateral sides of the epithelium. The morphology of immune cells was, however, different as compared to adult stages. Cells were generally less ramified and did not extend processes on epithelial cells or blood vessels (SuppleFig.3D2). Using acetyl-tubulin staining, we observed cilia in the tela choroidea, forebrain ChP and hindbrain ChP, but we could not detect axonal projections near blood vessels or epithelial cells (SuppleFig.3D1). Taken together, our results suggests that while all cellular components of the tela choroidea-ChPs are present at juvenile stages, the ChP is not yet fully mature, especially regarding the morphology of immune cells and axonal innervations.

### Expression of transporter genes involved in CSF secretion is evolutionarily conserved across vertebrates

Prior work from several laboratories pinpointed a critical set of ChP transporters involved in CSF secretion in mammals (Damkier *et al*, 2013; MacAulay *et al*., 2022; Oernbo *et al*, 2022). To identify whether these transporters are expressed in the zebrafish ChP, we extracted total RNAs from dissected adult forebrain ChP and performed bulk total RNA sequencing. We then compared our zebrafish transcriptomic data with publicly available human, rat and mouse ChP data (SuppleFig.4 and Fig.2) (Andreassen *et al*, 2022; Planques *et al*, 2021; Rodriguez-Lorenzo *et al*, 2020). We found 12,230 genes expressed in the ChP which were conserved across all 4 species, corresponding to 65-74 % of all protein coding genes (Fig.2A and Supplementary Fig.4A, C, D). Notably, 13-118 genes show species-specific expression, and 314 were mammalian specific (Fig.2A).

Next, we evaluated the expression of transporters and identified 576 conserved genes (Fig.2B and Supplementary Fig.4E). We found that the expression levels of the transporters were generally similar across species, with few exceptions such as SEC61G, SLC4A5 or TRPM3 which showed higher or lower ranks across species (Fig.2C). We next verified whether the transporters proposed to be involved in CSF secretion in mammals are also present in the zebrafish ChP (Fig.2D). Remarkably, we identified that all 7 CSF transporters are expressed in the zebrafish ChP although expression levels of AQP1 and SLC4A10 were lower, and SLC4A7 was higher in the zebrafish ChP (Fig.2D) compared to the mammalian ChPs. Altogether, our results show that the expression of CSF transporters is evolutionarily conserved across vertebrates and that the zebrafish ChP thus has the potential to secrete CSF.

### Generation of a novel choroid plexus specific driver line shows that the choroid plexus epithelial cells secrete proteins into CSF

*Fabp7b* (*fatty acid binding protein 7b*), which encodes for brain lipid binding protein (BLBP), is strongly expressed in the choroid plexus based on our transcriptomic data (*fabp7b*, TPM: 2627.58) and a prior report (Liu *et al*, 2003). To identify whether *fabp7b* could be a good marker for the choroid plexus, we performed *in situ* hybridization at 7 dpf and confirmed that it is specifically and strongly expressed in the forebrain and hindbrain ChPs, as well as in a subset of cells of the tela choroidea (Fig.3A, black arrow) and in the subcommisural organ, yet at lower levels (Fig.3A, white arrow).

Given the specific expression of *fabp7b* in the choroid plexus, we decided to create a knock-in Gal4 driver line to label and manipulate the ChP. Using the short homology mediated strategy (Welker *et al*, 2021), we inserted a Gal4 cassette in the exon 4 of *fabp7b* gene and generated the *Gt(fabp7b:2A-Gal4vp16)*(Fig.3B1 and SuppleFig.5A). To verify the specificity of our Gal4 knock-in line, we crossed it with the UAS effector line *Tg(uas:NTR-mCherry)*. We observed high expression of mCherry in the ChPs and some in the tela choroidea and subcommisural organ, which recapitulated our RNA expression pattern (Fig.3B2 and SuppleFig.5B). Fortunately, we did not observe mCherry signals in other cell types of the larvae (SuppleFig.5B). Additionally, we confirmed that BLBP antibody recognizes fabp7b in the tela choroidea-ChPs (Supplementary Fig.5C, white arrows) as well as fabp7a in radial glial cells (Adolf *et al*, 2006).

Next, we checked the coverage of the ChPs and tela choroidea by the *Gt(fabp7b:2A-Gal4vp16);Tg(uas:NTR-mCherry)* line, by crossing it with the *Gt(foxj1b:GFP)* line, which also labels the epithelial cells in the ChP and tela choroidea (Tian *et al*., 2009). We found that *Gt(fabp7b:2A-Gal4vp16)* covers 60-75% of the ChP (SuppleFig.5D, 5E) and only 25% of the tela choroidea (SuppleFig.5D, 5E) at 17 dpf, thereby allowing us to label and manipulate mainly the epithelial cells of the ChPs.

Using this ChP specific line, we tested whether epithelial cells of the ChPs have the potential to secrete proteins into CSF. To do so, we generated a transgenic line, *Tg(uas:starmaker-GFP)* which expresses starmaker, a 66 kDa secreted protein usually expressed in the otic vesicle (Sollner *et al*, 2003), fused with GFP. By crossing the *Gt(fabp7b:2A-Gal4);Tg(uas:NTR-mCherry)* line with *Tg(uas:starmaker-GFP)* line, we observed starmaker-GFP signals throughout the ventricles at 17-18 dpf (Fig.3C1-3C5). Notably, we did not detect signals elsewhere in the brain, suggesting that the ChP epithelial cells secrete proteins exclusively into the ventricle via their apical surfaces.

### Ablation of the choroid plexus epithelial cells decreases the size of the brain ventricles

Using our novel transgenic line, we aimed to investigate the role of ChP epithelial cells in CSF production and CSF-blood barrier. To this end, we focused our analysis on 17-18 dpf juveniles based on our observations that the ChP is relatively mature, and that the brain ventricular system is expanding at this stage (D’Gama *et al*., 2021). To label CSF, we developed an experimental strategy based on the observation that fluorescent dyes leak into the ventricles following cardinal vein injection (SuppleFig.3A1, 3C1). First, we confirmed that the intravenous injection really labels the brain ventricles by injecting the 10 kDa dextran Alexa-647 into *Gt(fabp7b:2A-Gal4vp16);Tg(uas:NTR-mCherry);Tg(uas:starmaker-GFP)* which secretes GFP-labelled starmaker in CSF (SuppleFig.6A and SuppleVideo1). Confocal imaging done 1-2 hours after injection in 17-18 dpf revealed a clear colocalization of the fluorescent dye with starmaker-GFP proteins throughout the ventricular system (SuppleFig.6A2-6A3), confirming the validity of our approach. Importantly, this method allows us to measure both the ventricle size and the integrity of the blood-CSF barrier in a less invasive way as compared to a conventional ventricular injection (Gutzman & Sive, 2009; Olstad *et al*., 2019).

To investigate the role of ChP epithelial cells in ventricle homeostasis, we chemogenetically ablated *fabp7b*- expressing cells by the metronidazole (MTZ)- nitroreductase (NTR) system (Curado *et al*, 2008) using the *Gt(fabp7b:2A-Gal4vp16);Tg(uas:NTR-mCherry);Gt(foxj1b:GFP)* line and its sibling controls (Fig.4A). We treated 14 dpf juveniles with DMSO or MTZ for 24 hours. Following a recovery of 2-3 days, we injected the 10 kDa dextran Alexa-647 into the cardinal vein of the fish and imaged them after 1-2 hours (Fig.4A). First, we verified the ablation efficiency after MTZ treatment (Fig.4B-C), by measuring the surface area of the tela choroidea-ChP, which expresses *foxj1b*-GFP. We observed a reduction of tela choroidea and ChP size by 70-77% in the ablated fish compared to our two control groups: DMSO treated NTR-mCherry positive fish and MTZ treated NTR-mCherry negative fish (Fig.4C-D). Interestingly, we noticed that the MTZ ablation showed similar yields in the tela choroidea as compared to the ChP although only 25% of the tela choroidea expressed NTR-mCherry on average (SuppleFig.5E). Next, we measured the size of the brain ventricles following ablation (Fig.4E-G). We found that all brain ventricles were strikingly smaller in the ablated fish, compared to the controls (Fig.4F). To quantify ventricular sizes, we measured the areas of the telencephalic, di-/mesencephalic and rhombencephalic ventricles (Fig.4E, F). We found that the ventricles in the ablated fish were reduced by circa 5-40% as compared to controls (Fig.4G). Notably, we did not observe any differences in fish size and brain size between control and ablated fish (SuppleFig.7). Intriguingly, we did not detect notable difference of fluorescent intensities neither in CSF nor in the brain parenchyma between control and tela choroidea-ChP ablated fish (SuppleFig.8A-8D), suggesting that the brain barriers were unaffected by our manipulation. Altogether, our findings identify a major role of the ChP in maintaining ventricular volume, most probably through its role in CSF secretion (Fig.4H).

**Figure 4.**
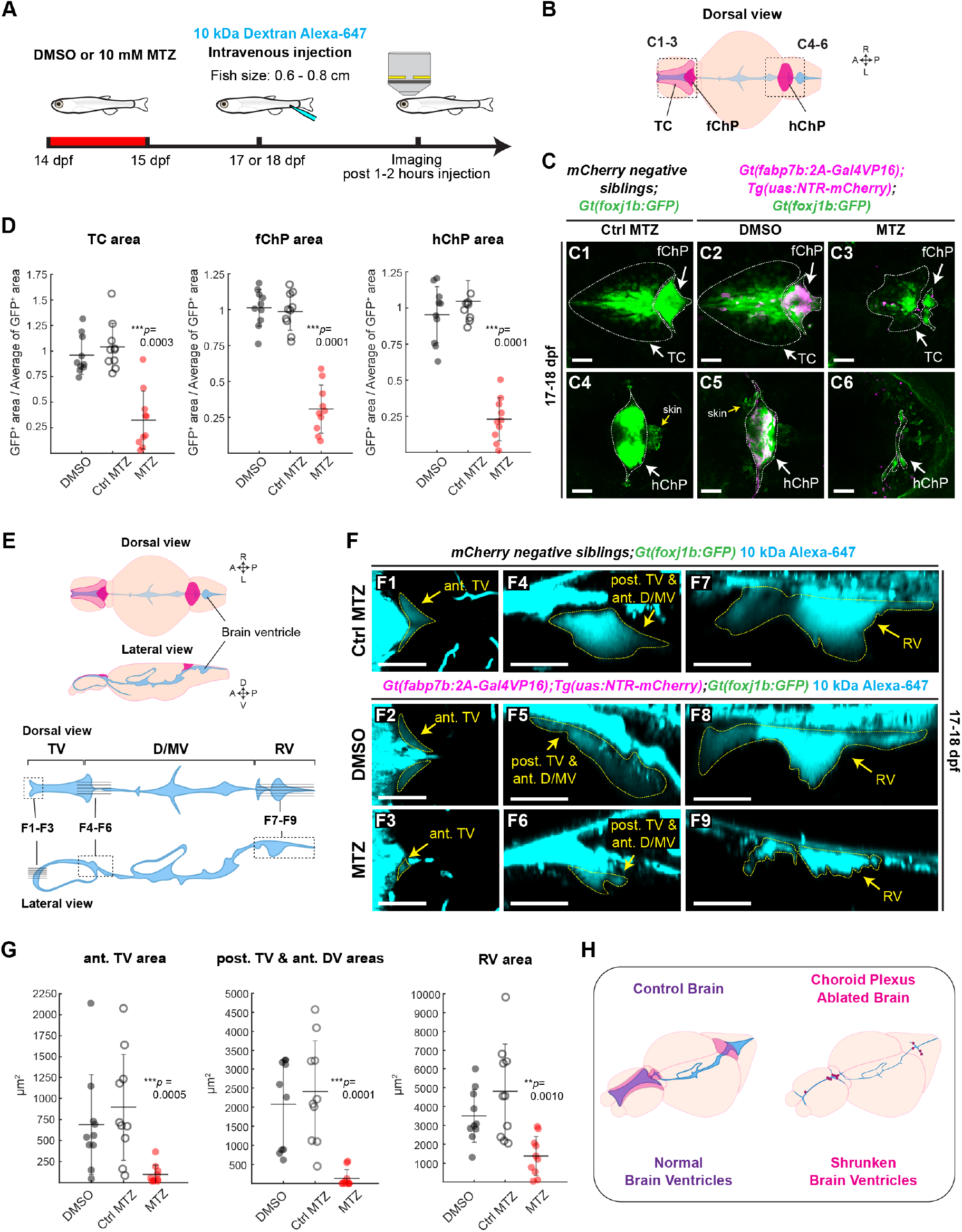
The zebrafish choroid plexuses are required for the maintenance of brain ventricles. **(A)** Experimental scheme used for measuring brain ventricular size and blood-CSF barrier integrity following MTZ-mediated ablation and injection of a 10 kDa dextran Alexa-647 fluorescent dye in the cardinal vein. **(B)** Diagram of the juvenile brain parenchyma and ventricles with insets indicating the location of panels (B1-B6). **(C)** Max projection images of the TC, fChP and hChP in the control fish (C1-C2, C4-C5) and TC-ChPs ablated fish (C3, C6) at 17-18 dpf. **(D)** Quantification of the TC, fChP and hChP areas in the control fish (DMSO and Ctrl MTZ) and TC-ChPs ablated fish (MTZ). *n = 10.* The TC and ChPs size in the ablated fish are shown as normalized GFP^+^ ratio (mean+/-STD): TC, 0.32+/-0.28; fChP, 0.31+/-0.17; hChP, 0.23+/-0.15. **(E)** Diagrams of dorsal and lateral views in the juvenile brain (upper) and the brain ventricles (lower). Projections were made of 6 horizontal slices in the dotted boxed F1-F3, 5 sagittal slices in the F4-F6, and 5 sagittal slices in the F7-F9, as indicated in the scheme. **(F)** Horizontal **(F1-3)** and sagittal **(F4-F6, F7-F9)** views of the ventricles following a 10 kDa Alexa-647 injection in control **(F1-F2, F4-F5, F7-F8)** and TC-ChPs ablated fish **(F3, F6, F9)** at 17-18 dpf. Ventricles are highlighted with dotted yellow lines. **(G)** Quantification of the ant. TV, post. TV & ant. D/MV and RV area in the control fish (DMSO and Ctrl MTZ) and TC-ChPs ablated fish (MTZ). n = 10. **(H)** Diagrams of the brains in the control and TC-ChPs ablated fish. Statistical analysis: Kruskal-Wallis test. \*\**p* < 0.01, \*\*\**p* < 0.001. Abbreviations: TC, tela choroidea; fChP, forebrain choroid plexus; hChP, hindbrain choroid plexus; ant., anterior; post., posterior; TV, telencephalic ventricle; D/MV, dien-/mesencephalic ventricle; RV, rhombencephalic ventricle. A, anterior; P, posterior; D, dorsal; V, ventral; R, right; L, left; STD, standard deviation. Scale bars: 50 µm

## Discussion

By performing histological, molecular, and physiological analyses of the zebrafish ChP, our study highlights the evolutionary conservation of the ChP in vertebrates. Using novel genetic tools, we also provide a deeper mechanistic understanding on the function of ChP in maintaining ventricular homeostasis. Due to conserved features of the zebrafish brain and brain barriers, we trust that our tools in the zebrafish will be powerful to better understand the ChP function in the brain.

In mammals, each ventricle has one choroid plexus, with a total of four. In zebrafish, there has, only been two ChPs described so far based on anatomical and gene expression profiles, which are located in the forebrain and hindbrain (Garcia-Lecea *et al*., 2008; Korzh, 2023). Given that the forebrain ChP is located between the pineal gland and the habenula and is nearby the subcommisural organ (Korzh, 2023), it most probably corresponds to the ChP of the 3^rd^ ventricle in mammals. Interestingly, the forebrain ChP, but not the hindbrain ChP, folds into a complex 3D structure between the juvenile and adult stage, when the forebrain ChP expands steadily. At this point it is not clear why only the forebrain ChP changes its shape dramatically and not the hindbrain ChP. One possibility would be that by generating these folds it increases the epithelial surface area and thereby can secrete more CSF in a relatively confined area. However, it was shown that the most active ChP in mammals is the one located in the 4^th^ ventricle and not the 3^rd^ (Gomez & Potts, 1981; Keep *et al*, 1986; MacAulay *et al*., 2022; Quay, 1966). Future work identifying the contribution of individual ChP in CSF dynamics are required to resolve these questions. Besides, it remains unclear why zebrafish do not have a ChP in the telencephalon. We show here that the tela choroidea has similar anatomical features as the ChP, containing epithelial cells that harbour brushes of microvilli and that are connected with tight junctions. Originally the tela choroidea refers to a primordium of the ChP in the developing mammalian brain. However, it has also been used to designate a thin ventricular layer that cover the dorsal telencephalon in ray-finned fishes (Folgueira *et al*, 2012; Nieuwenhuys, 2011; Porter & Mueller, 2020) and contain motile ciliated cell lineages (D’Gama *et al*., 2021). Based on our data, we suggest that the tela choroidea may have ChP-like functions beside playing a role as an ependymal cell layer.

Prior work in rodents using bulk and single cell transcriptomics revealed differences in gene expression between the different ChPs, especially the lateral ventricle and 4^th^ ventricle (Lun *et al*, 2015a), and during developmental stages (Dani *et al*., 2021). In our study, we investigated only the adult forebrain ChP given its amenability to be dissected out. However, it will be interesting to perform a more precise analysis of the transcriptome of the tela choroidea and ChP throughout development. We trust that this information will allow to better identify the mammalian counterpart of each zebrafish ChP for future work exploring the exact roles of the tela choroidea and ChPs in brain physiology.

In our transcriptomic analyses, we identified that the transcriptomes of the vertebrate ChPs are well conserved. The transporters proposed to be involved in CSF secretion in mammals are highly expressed also in zebrafish, which underscores the CSF-secreting ability of zebrafish ChP. Yet, we noticed that the expression levels of some genes are different between species, including TTR (*transthyretin*), which is very lowly expressed in the zebrafish ChP (TPM: zebrafish, 0.36) as compared to mammals (TPM: human, mouse, rat > 239,000). Notably, the expression of some transporter genes was slightly different, which may result in distinct ion levels or solute content in CSF across species. The precise ion concentration in zebrafish CSF is currently unknown, yet artificial CSF used in laboratories are generally slightly different for mammals and fish (2017; Jeong *et al*, 2022). Future comparative work on CSF properties across species could reveal fundamental new knowledge on the mechanisms of brain homeostasis.

Developing specific genetic tools to manipulate the ChP in a time-specific manner is a game changer. It has been so far very difficult to specifically target the ChP using genetics, and often researchers have made use of the AAV2/5 serotype which preferentially target the ChP epithelial cells in rodents (Chen *et al*, 2020; Haddad *et al*, 2013; Jang & Lehtinen, 2022; Sadegh *et al*, 2023; Xu *et al*, 2021). Here we generated a versatile genetic tool using the Gal4-UAS system allowing for the first time to specifically investigate and manipulate the ChP epithelial cells in the zebrafish. *Fabp7b*, which encodes BLBP, is mainly expressed in the ChP, while its orthologue *fabp7a* is expressed in radial glia cells (Adolf *et al*., 2006). Based our transcriptomics analysis, it appears that *Fabp7* is lowly expressed in mammalian ChP (TPM: human, 3.37; rat, 0.51; mouse; 1.12). Also, given its expression in neuronal progenitors (Anthony *et al*, 2004; Kurtz *et al*, 1994), fabp7 would probably not serve as a good marker for the mammalian ChP.

Using these tools, we were able to not only chemogenetically ablate ChP epithelial cells, but also start investigating CSF dynamics *in vivo* in an intact brain. Notably, we were successful at labelling the brain ventricles with a fluorescent protein using our newly created *Tg(uas:starmaker-GFP)* line. Currently, most studies use injection of fluorescent dyes or proteins into the ventricles to visualize CSF or the brain ventricles in mammals and zebrafish (Fame *et al*, 2016; Olstad *et al*., 2019; Sweeney *et al*, 2019). This is, however, an invasive way that can disturb the ventricular system, by inducing wound-driven inflammation, and changing the ventricle volume, content and osmolarity due to the addition of exogenous molecules. By offering the possibility to measure the morphology and size of the brain ventricles in a non-invasive manner, we are confident that our versatile tool opens new possibilities to study CSF-associated diseases like hydrocephalus.

In our study, we showed that chemogenetic ablation of the ChP epithelial cells reduces the size of the brain ventricles dramatically. We suggest that reduced CSF secretion due to the loss of epithelial cells underlies the shrunken ventricles, and thus supports a pivotal role for the ChP in CSF secretion (MacAulay *et al*., 2022; MacAulay & Toft-Bertelsen, 2023). Based on prior work, it is expected that the amount of CSF and thereby the volume of the brain ventricles is maintained through a balanced CSF secretion and clearance (Bothwell *et al*., 2019). Our data do not however align with the proposed role for the ChP as CSF drainage pathway (Sadegh *et al*., 2023; Xu *et al*., 2021), at least at the tested age group and experimental conditions. By reducing CSF production in ChP ablated fish, this would lead to a disturbed balance resulting in a loss of the hydrostatic pressure necessary to maintain the ventricular sizes. There has been reports of extrachoroidal CSF production (Curl & Pollay, 1968; Pollay & Curl, 1967). Given the massive reduction of brain ventricles in our ChP ablated fish, the contribution of these sources may be minimal at the investigated age. It is also possible that the reduced surface area of the tela choroidea-ChPs following cell loss push down on the ventricles and contribute to their smaller size. Alternatively, CSF may leak out from the ventricles due to disrupted barriers. We have however not observed any changes in the intensity of intravenously injected dyes within the ventricles or parenchyma, suggesting that at least 2-3 days after ablation the brain barriers are not impaired. To date, CSF clearance pathways are poorly characterized in zebrafish. Hence it is not possible to investigate whether CSF clearance is affected upon ChP ablation. Taken together, our experiments provide a valid proof that the ChPs are involved in CSF secretion, and thereby maintain ventricular volume.

It is very puzzling that despite a loss of circa 70 % of the epithelial cells in the ChPs, we did not observe a disruption of the CSF-blood brain barrier. There are multiple possibilities to explain the phenomenon. Cell extrusion, which is a mechanism extruding damaged cells out of epithelial tissues undergoing apoptosis (Guillot & Lecuit, 2013), could keep the integrity of the epithelium and sustain CSF-blood barrier function. It is however hard to conceive that the remaining 30 % epithelial cells could cover all the surface of the damage epithelium and maintain the barrier. There might be additional mechanisms involved such as wound healing and regeneration given that we waited for 2-3 days prior to performing the intravenous injection. In our study, we used a 10 kDa dextran-based fluorescent dye. It is possible that other molecules, with a smaller size or different structure, could show different penetrance to the ventricles. To date, it is not even clear to us how the dye penetrates the ventricle in the control conditions. An earlier study identified that both the blood-brain and the blood-CSF barriers were more permeable upon loss of the tight junction claudin 5 in zebrafish (Li *et al*, 2022). Due to very low fluorescence intensities in the parenchyma, we do not expect it to go through the blood-brain barrier. Instead, it may go through the leaky circumventricular organs, such as the pituitary, which have fenestrated capillaries (Gordon *et al*, 2019). Whether it goes in the CSF through the ChP or the circumventricular organs will, however, require future investigations. Notably, *fabp7b* is also expressed in the subcommisural organ, a circumventricular organ secreting SCO-spondin, the main component of the Reissner’s fiber that is important for spine morphogenesis in zebrafish (Rose *et al*, 2020; Troutwine *et al*, 2020). A recent preprint identified that SCO-spondin knockout mice has small brain ventricles with mild spine deformation (Xu *et al*, 2023), suggesting that the subcommisural organ might also influence the brain ventricle size. Future work on the subcommisural organ is required to identify how it maintains the brain ventricle size and whether it could contribute to the phenotype we observed in the ChP ablated zebrafish.

Altogether, our work provides strong evidence that the ChP and brain ventricular system are highly conserved across vertebrates, and that the ChP serves to secrete CSF also in the zebrafish. Our findings thus position the zebrafish as a suitable animal model for disease modelling. By contributing versatile genetic tools in the zebrafish, we are confident that our work opens new avenues for unravelling novel insights on the impact of ChP and CSF on brain development and physiology in health and disease.

## Acknowledgements

We thank the Fluorescent Reporter Zebrafish Cooperation Center (FRZCC), Republic of Korea, for providing the zebrafish line (*FRZCC1104*), Valérie Wittamer (Université Libre de Bruxelles, Belgium) for sharing the 4C4 antibody, Sebastian Gerety (Wellcome Sanger Institute) for sharing the the pUAS-Self vector, the fish facility support staff in our respective institutes for zebrafish maintenance, and all members of the Jurisch-Yaksi and Yaksi laboratories for their feedback on this work. BioRender was used to generate some illustrations. This work was supported by a FRIPRO research grant from The Research Council of Norway (NJY 314189) to NJY, the National Research Foundation of Korea (NRF) grant funded by the Korea government (MSIT) (2022R1A2C1091336) to HCP, and the Lundbeck Foundation (to NM; R276-2018-403) the Fondation ARC pour la Recherche sur le Cancer (PJA2018208167 to MF, A2011 Postdoctoral fellowship to MP) and the Human Frontiers Science Program (CDA00036/2010 to MF).

## Author contributions

Conceptualization: NJY, IJ, NM, SNA, MF; Methodology: IJ, SNA, LH, MP, YS, MF; Formal analysis: IJ, NJY, SNA Investigation: IJ, SNA, NJY, LH; Resources: NJY, HCP, MF, NM ; Data curation: IY, NJY, SNA; Writing - original draft: IJ, NJY; Writing- review & editing: all authors; Visualization: IJ, NJY, SNA; Supervision: NJY, HCP, NM, MF; Funding acquisition: NJY, HCP, NM, MF, MP

## STAR Methods

### Key resources table

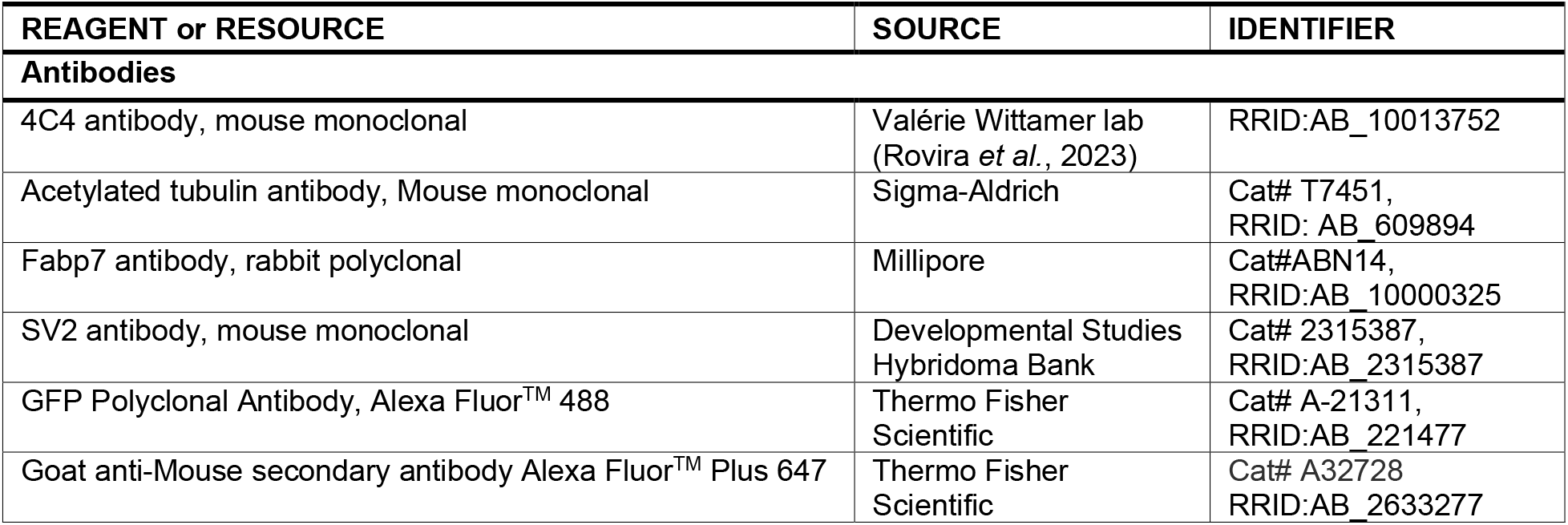
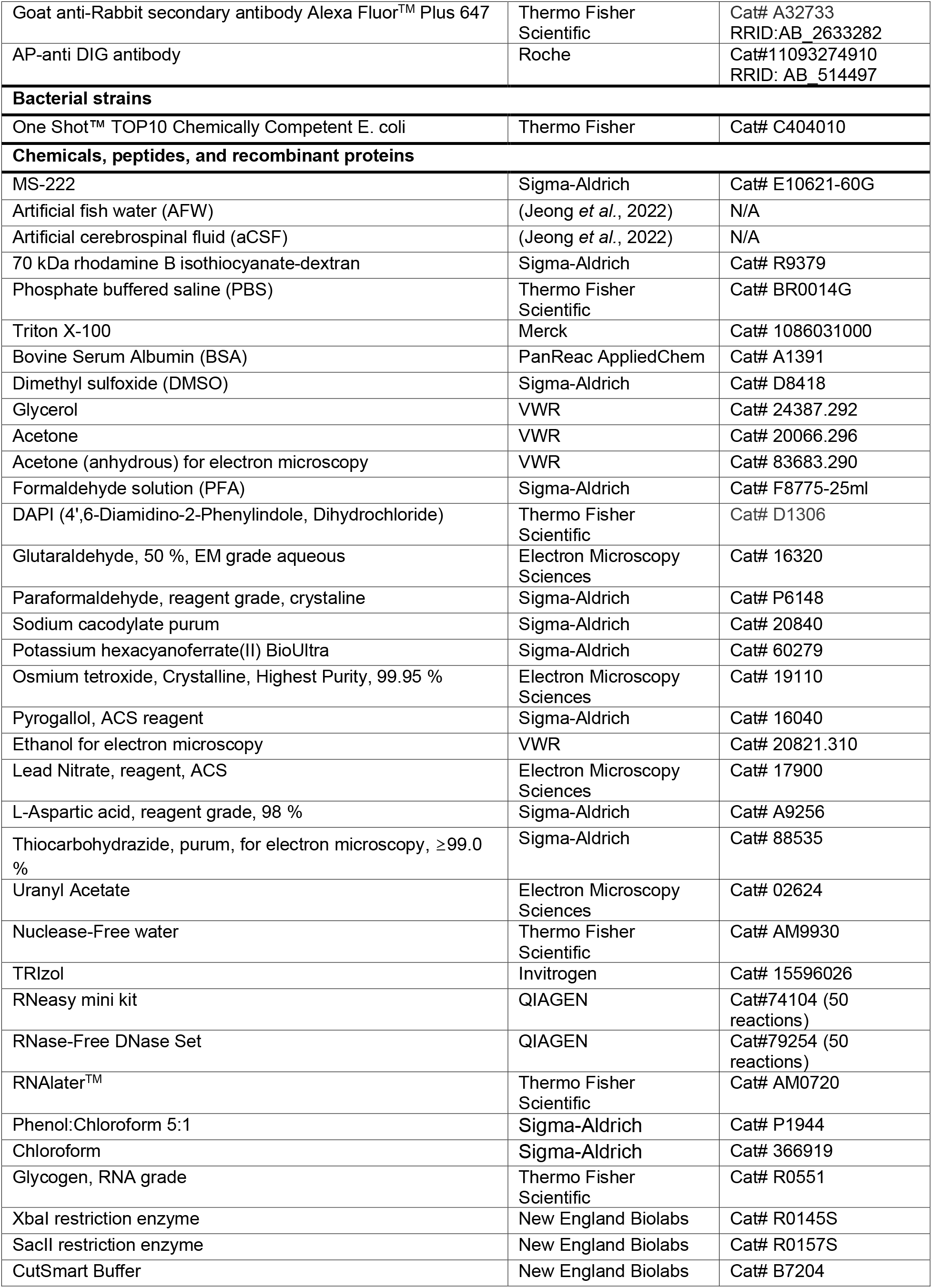
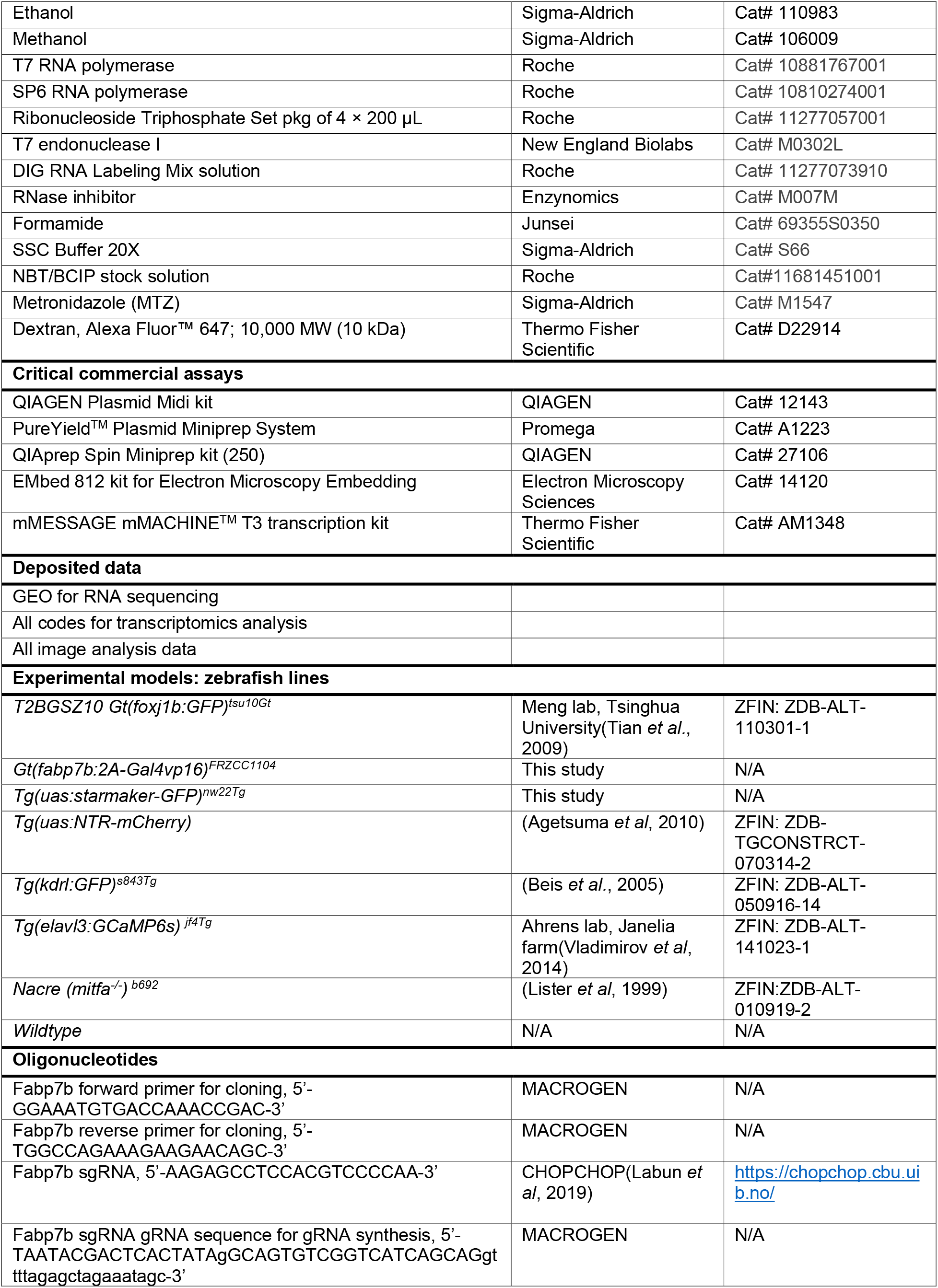
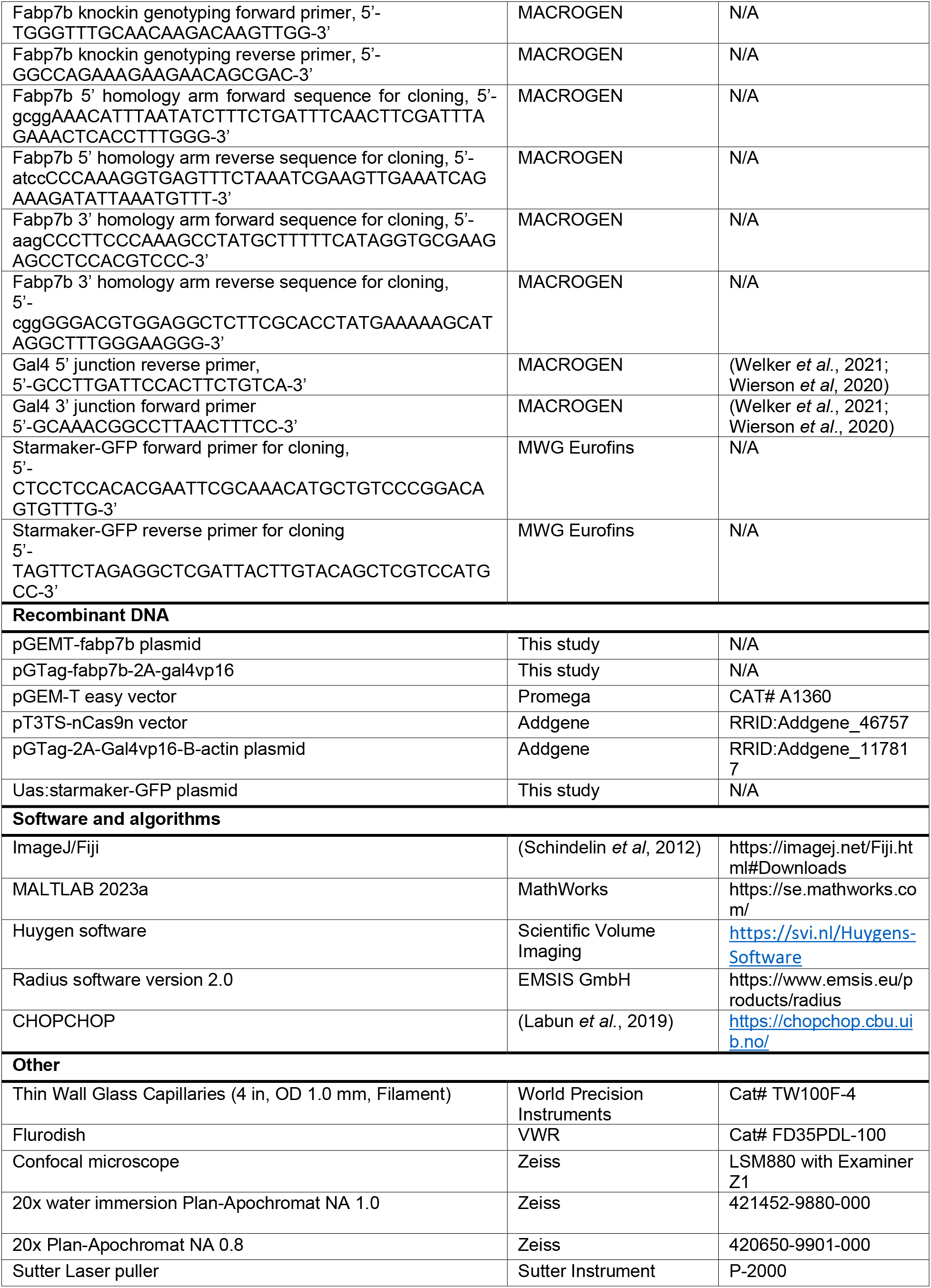
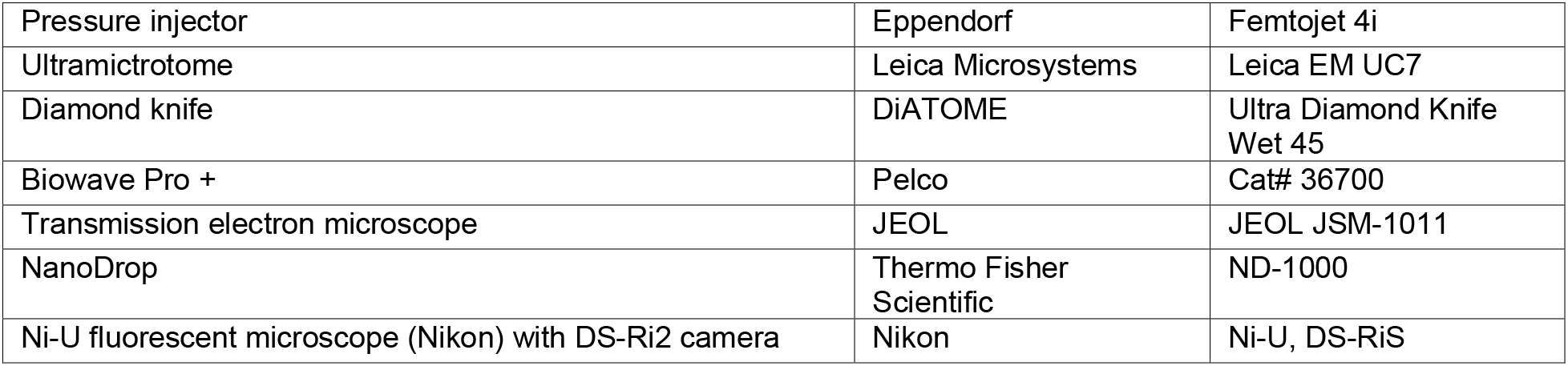

### Lead contact

Further information and requests for resources and reagents should be directed to and will be fulfilled by the lead contact, Nathalie Jurisch-Yaksi.

### Materials availability

Material generated in this paper will be shared by their creators (I.J., H.C.P: *Gt(fabp7b:2A- Gal4vp16)^FRZCC1104^*, I.J., N.Y.J: *Tg(uas:starmaker-GFP)^nw22Tg^*, M.F UAS:starmaker-GFP plasmid) upon request and may require completion of a material transfer agreement.

### Data and code availability

Raw imaging data used to generate the figures of the manuscript will be deposited upon publication. The RNA sequencing data will be available on GEO upon publication. Codes for the analysis of RNA sequencing are available at https://github.com/Sorennorge/Choroid-plexus-orthology-study

### Experimental model and study participant details

#### Zebrafish maintenance and strains

The animal facilities and maintenance of zebrafish, Danio rerio, were approved by the Norwegian Food Safety Authority and the Korea Disease Control and Prevention Agency. All the procedures were performed on zebrafish at different developmental stages post fertilization in accordance with the European Communities Council Directive, the Norwegian Food Safety Authorities (FOTS permit #28918: Roles of the choroid plexus in brain physiology), and the animal experiment guidelines of Korea National Veterinary Research and Quarantine Service. The experimental procedures performed in the Republic of Korea were approved by the Korea University Institutional Animal Care & Use Committee (IACUC; KOREA-2019–0165) and were performed in accordance with the animal experiment guidelines of Korea National Veterinary Research and Quarantine Service. Embryonic, larval, juvenile and adult zebrafish were reared according to standard procedures of husbandry at 28.5°C.

For our experiments, the following zebrafish lines were used: *Gt(foxj1b:GFP)* (Tian *et al*., 2009), *Tg(uas:NTR-mCherry)* (Agetsuma et al., 2010), *Tg(kdrl:GFP)* (Beis et al., 2005), *Tg(elavl3:GCaMP6s)* (Vladimirov et al., 2014), *Gt(fabp7b:2A-Gal4vp16)^FRZCC1104^* and *Tg(uas:starmaker-GFP)^nw22Tg^*). Zebrafish were analyzed irrespective of their gender. Note that for zebrafish younger than 2-3 months, gender is not yet apparent. All zebrafish lines containing the *Gt(fabp7b:2A-Gal4vp16)^FRZCC1104^* allele were maintained as heterozygous to avoid loss of fabp7b. Animals were either in the AB or pigmentless nacre*^b692^* (*mitfa*^−/−^) (Lister *et al*., 1999) background.

#### Generation of *Gt(fabp7b:2A-Gal4vp16)^FRZC1104^* knock-in zebrafish

To create the genetic line to label *fabp7b*-expressing cells specifically, the short homology mediated knock-in strategy was used (Rayamajhi *et al*, 2023; Welker *et al*., 2021; Wierson *et al*., 2020) First, a Cas9 target site was selected in exon 4 of *fabp7b* gene using by CHOPCHOP (http://chopchop.cbu.uib.no/). The sgRNA was synthesized by the cloning-free gRNA *in vitro* synthesis (Welker *et al*., 2021; Wierson *et al*., 2020) and Cas9 mRNA from the pT3TS-nCas9n plasmid (Addgene) with mMESSAGE mMACHINE^TM^ T3 transcription kit (Thermo Fisher Scientific). To validate efficiency of the sgRNA, *fabp7b* sgRNA and Cas9 mRNA was microinjected into one-cell wildtype embryos and genomic DNAs were extracted from circa 20 injected embryos at 24 hpf. Then, T7 endonuclease I assay was performed (Jao *et al*, 2013; Jeong *et al*, 2021). Next, homology arms of the target vector were designed by the GTagHD (http://www.genesculpt.org/gtaghd/) and the pGTag-2A-Gal4vp16-B-actin plasmid (Addgene) was used as a donor vector (Welker *et al*., 2021; Wierson *et al*., 2020). The *fabp7b* targeting donor plasmid including 2A-Gal4vp16-B-actin was constructed and the plasmid was purified with the QIAGEN Plasmid Midi kit (QIAGEN) and PureYield Plasmid Miniprep System (Promega). 2 nL of a mixture of *fabp7b* sgRNA (50 pg), universal gRNA (25 pg), Cas9 mRNA (200 pg) and the target donor plasmid (10 pg) was injected into one-cell stage embryos of several UAS transgenic lines. The injected embryos expressing fluorescence in the choroid plexus were screened and raised to adulthood (F0). To identify germline-transmitted lines, the F0 fish were crossed with several UAS transgenic lines and were validated based on choroid plexus specific fluorescence in F1 embryos. Following selection of the germline-transmitting founder, we confirmed the integration of the 2A-Gal4vp16-B-actin DNA into the genome by PCR on the genomic DNAs of the fluorescence positive larvae. We verified that the 2A-Gal4vp16 DNA was integrated into the genomic area by performing PCR and sequencing at the 5’ and 3’ junctions of integration sites (SuppleFig.5A).

#### Generation of Tg(uas:starmaker-GFP)^nw22Tg^ line

The open reading frame (ORF) of *starmaker* (*stm*: ENSDARG00000035694) was amplified by PCR from zebrafish cDNA. The amplified PCR products were fused to a C-terminal eGFP-Tag separated by a 5’-TCTAGAACTATA-3’ linker sequence containing an XbaI restriction enzyme site. The starmaker-GFP DNA was inserted into the pUAS-Self vector (gift from Sebastian Gerety) downstream of the zebrafish-optimized 5’-GCAAAC-3’ Kozak sequence through Gibson cloning. The starmaker sequence in this plasmid construct lacks the 45 base pairs corresponding to Exon 15 of the Ensembl transcript stm-202 (ENSDART00000105174.6), resulting in a 15 amino acid in-frame deletion.

To generate the transgenic line, 2 nL of a mixture of the uas:starmaker-GFP plasmid DNA (60 pg) and tol2 mRNA (10 pg) was microinjected into one-cell stage embryos. The injected embryos expressing red fluorescence in the eye lens were selected and raised to adulthood (F0). Germline-transmitting founders were identified by breeding with several Gal4 transgenic lines and selected based on RFP expression in the lens and predicted GFP signals in the Gal4 lines. Stable F1 embryos showing strong RFP signals were raised to adult zebrafish.

### Method details

#### Immunohistochemistry and confocal microscope imaging

Zebrafish were euthanized by cold artificial fish water (AFW) or high dose MS222 in AFW. Larvae and juvenile were fixed in 4% paraformaldehyde (PFA), 1% DMSO and 0.3% Triton X-100 in PBS (phosphate buffered saline) (0.3% PBST) solution at 4°C overnight. Then, the fixed samples were washing with 0.3% PBST three times and then the brains were dissected out. For adult, the brains were dissected out in cold artificial cerebrospinal fluid (aCSF) and then fixed with 4% PFA at 4°C overnight. The pineal gland was removed in samples used to image the forebrain choroid plexus (ChP).

The samples were washed with 0.3% PBST three times for 30 min and then permeabilized by −20°C stored acetone for 10 min (larvae), 15 min (juvenile) and 1 hour (adult) at room temperature (RT). Then the brains were washed again with 0.3% PBST three times for 30 min and immersed with blocking solution (0.1% bovine serum albumin (BSA) and 1% DMSO in 0.3% PBST) for over 2 hours on shaker at RT. Samples were incubated in the blocking solution with anti-acetylated tubulin (1:500, Sigma-Aldrich), anti-4C4 (1:400, gift from Wittamer lab) (Rovira *et al*., 2023), anti-Fabp7 (1:400, Sigma) and anti-SV2 (1:100, Developmental Studies and Hybridoma bank) antibodies overnight at 4°C. On the next day, samples were washed with 0.3% PBST three times for 3 hours and then immersed in the blocking solution for 2 hours on a shaker at RT. Subsequently, samples were incubated at 4°C overnight with secondary antibodies (goat anti-rabbit Alexa Fluor^TM^ Plus 647 or goat anti-mouse Alexa Fluor^TM^ Plus 647 (Thermo Fisher Scientific, 1:1000) and Alexa Fluor 488 tagged anti-GFP polyclonal antibody (Thermo Fisher Scientific, 1:500) to boost the GFP signals. On the third day, samples were incubated with 0.1% DAPI (Thermo Fisher Scientific) at RT for 30 min on shaker and then washed with 0.3% PBST three times for 3 hours on shaker. Next, samples were transferred to the PBS solutions gradually increased glycerol concentrations (25%, 50% and 75% glycerol in PBS). After staining steps were finished, samples were imaged immediately or stored in 75% glycerol in PBS at 4°C before imaging. Samples were imaged using by a Zeiss LSM880 confocal microscope with an Examiner Z1 and a 20x Plan NA 0.8 objective.

#### Brain ventricle injection and confocal microscope imaging

To visualize the brain ventricles and CSF, we followed our previous protocols (D’Gama *et al*., 2021; Jeong *et al*., 2022). Adult zebrafish were first euthanized by cold AFW. The adult brains were then dissected in cold aCSF and mounted on sylgard using a metal pin (Jeong *et al*., 2022; Kermen *et al*, 2020a; Kermen *et al*, 2020b). The injection mixture containing 70 kDa rhodamine B isothiocyanate-dextran (RITC-dextran; Sigma-Aldrich) dissolved in aCSF at a final concentration of 10 mg/mL. The injection needles were pulled with a Sutter Instrument Co. Model P-2000, from thin-walled glass capillaries (1.00 mm; WPI). The needle tip was opened by cutting with fine forceps. A microinjector (Eppendorf Femtojet 4i) was used to microinject around 15 nL of the mixture solution into the telencephalic ventricle. After injection, the adult brain explants were instantly transferred to the confocal microscope and imaged with a 20x water-immersion objective (Zeiss, NA 1.0, Plan-Apochromat) at RT. The laser power was adjusted according to the depth of imaging.

### Transmission electron microscopy

#### Sample fixation, staining and embedding

Dissected adult brains were trimmed nearby the forebrain ChP by a razor blade in cold aCSF, and fixed with 2.5% glutaraldehyde and 4% PFA in 0.15 M cacodylate (pH 7.4) overnight at RT. For the dissection, the pineal gland was not removed and the dissected brains were trimmed into smaller pieces. Next, the trimmed brains were encapsulated with 4% ultra-low melting agarose in 0.1M PB (phosphate buffer) and the access of agarose around the trimmed brain were removed with a razor blade. The samples were put in 2.5% glutaraldehyde in 0.15 M cacodylate (pH 7.4) and storage at 4°C. For preparation and *en bloc* staining a microwave (Pelco Biowave Pro+) was used with an established protocol with minor modifications (Hua *et al*, 2015; Mikula & Denk, 2015)(SuppleTable1). The procedure includes successive exposures of the sample to 2% OsO4 in 0,15 M cacodylate buffer, 2,5% potassium ferrocyanide in 0,15 M cacodylate buffer, filtered 320 mM pyrogallol in aqueous solution (pH 4.1), 1% OsO4 in aqueous solution, filtered 320 mM pyrogallol in aqueous solution (pH 4.1), 1% OsO4 in aqueous solution, filtered 1 % uranyl acetate in aqueous solution and filtered lead aspartate solution (0.02 M lead nitrate and 0.03 M aspartic acid, pH 5.5). Double rinses in pure filtered water were performed between the staining steps.

Samples were then dehydrated through a graded ethanol series (20%, 50%, 70%, 90%, 100%, 100%) and then into pure acetone (2 x 100%). After acetone, samples were infiltrated and incubated in 812 Epon resin diluted with ethanol (25%, 50%, 75%, 100%). In the end samples were placed in embedding molds with pure resin and cured at 60 °C for 75 h.

#### Sample sectioning

Embedded samples were trimmed using an ultramicrotome (Leica EM UC7) with a dry glass knife. Sections of 1 µm were cut, put on a microscope slide and stained with toluidine blue for orientation with light transmission microscopy. The area of interest was then selected for ultrathin sectioning. Ultrathin sections (60 nm) were cut with an ultramicrotomy (Leica EM UC7) using a diamond knife (Diatome), and subsequently collected on slot Copper grid with Formvar support film.

#### Microscopy

The grids were viewed with a TEM (JSM-1011, JEOL) operating at 80 kV. Images were captured with a Morada digital camera with Radius software version 2.0 (EMSIS GmbH).

### Total RNA sequencing and transcriptomic analysis

#### Total RNA preparation and RNA sequencing

To extract total RNA from the forebrain ChP, adult zebrafish (Nacre and AB, 6 - 8-month-old, > 3 cm) were euthanized by cold AFW and the brains were dissected out in cold aCSF. The forebrain ChPs were collected after removing the pineal gland. The dissected forebrain ChPs were kept temporarily in 1.5 mL tubes containing 20 μL RNAlater^®^ (Thermo Fisher Scientific) on ice until RNA extraction steps started. Two batches of samples were prepared, and each batch had from 12 to 19 forebrain ChPs. To lyse the forebrain ChPs, TRIzol solution (500 μL, Thermo Fisher Scientific) was added and the tissues were homogenized through 27-gauge needles 15-18 times. More TRIzol (500 μL) was added into the tubes and the dissociated tissues were incubated for 5 min at RT. Chloroform (Sigma-Aldrich, 200 μL) was added to the samples and the tubes were inverted for several times to mix them. These were kept for 2 min at RT and centrifuged at 4°C for 15 min at 12,000 rpm. Then colorless supernatants including total RNAs were purified by two different methods using glycogen, or spin columns.

For the glycogen method, the RNA supernatant was transferred to a new tube and 0.5 μL glycogen solution (20 mg/mL, final glycogen mass: 10 μg) was added. Then, isopropanol (500 μL) was added to the sample, mixed by inverting and incubated on ice for 10 min. Then, the sample was centrifuged for 15 min at 12,000 rpm at 4°C. The supernatant was discarded, and the pellet was washed with 1 mL 70% ethanol (w/nuclease-free water) chilled at −20°C. The pellet was air-dried on ice for 2-5 min and dissolved in 70 μL nuclease-free water (Invitrogen). The sample was then treated with 10 μL DNase enzyme (QIAGEN) for 10 min at RT to remove DNA in the RNA samples. To isolate the RNAs, phenol-chloroform ethanol precipitation was performed. The DNase treated RNA sample was added to phenol-chloroform (pH8.0, company name, 100 μL) and mixed for 5-10 sec by inverting. The RNA was centrifuged for 8 min at 12,000 rpm at 4°C. The clear supernatant was transferred to a new tube, treated with chloroform (100 μL) and mixed for 5-10 sec by inverting. Following centrifugation at 12,000 rpm for 8 min at 4°C, the supernatant was precipitated with 10 μL 4M Ammonium Acetate buffer (100 μL) and 100% ethanol 330 μL for 30 min at −20°C and centrifuged at 12,000 rpm for 30 min at 4°C. The pellet was washed with 70% EtOH (w/nuclease-free water, 500 μL) and air dried on ice for 2-5 min. The pellet of RNAs was dissolved in 20 μL nuclease-free water (Thermo Fisher Scientific).

For the spin column method, the RNA supernatant was mixed to identical amount of isopropanol, loaded on RNA grade spin columns (QIAGEN, RNeasy Mini kit) and centrifuged for 30 sec at 8,000 rpm. The spin columns were washed with 700 μL Buffer RW1 and 350 μL Buffer RW1 followed by DNase enzyme treatment in Buffer RDD for 10 min (10μL DNase in 70 μL Buffer RDD per column) at RT. After treatment, the column was washed with 350 μL Buffer RW1 and 500μL Buffer RPE. To elute the RNAs from the columns, 20 μL nuclease-free water was added and incubated for 2 min. The RNAs were centrifuged for 1 min at 8000 rpm and collected in tubes. The concentrations of four eluted RNAs were quantified using Nanodrop (Thermo Fisher Scientific) and stored at −80°C before RNA sequencing.

The quality of RNA samples was analysed by bioanalyzer at Novogene. Due to low amount of quantity in the RNA samples, the RNAs were pre-amplified using by the SMARTer kit and RNA sequencing (paired-end 150 bp, with 12 Gb output) was performed on an Illumina NovaSeq 6000 (Illumina) at Novogene.

#### Transcriptomic analysis

RNA sequencing data for human, rat, and mouse were obtained from the GEO repository (Barrett *et al*, 2013; Edgar *et al*, 2002). Rat choroid plexus transcriptome data was obtained with GEO accession number GSE194236 (sample: SRR17713531) (Andreassen *et al*., 2022). The human choroid plexus transcriptome data was obtained with GEO accession number GSE137619 (samples: SRR10134643-SRR10134648) (Rodriguez-Lorenzo *et al*., 2020). The mouse choroid plexus transcriptome data was obtained from GEO accession number GSE157386 (samples; SRR12576642-SRR12576645, SRR12576650-SRR12576653) (Planques *et al*., 2021). The zebrafish choroid plexus transcriptome data was obtained from this study.

All program parameter settings for library building and mapping, together with all scripts for the gene annotation and analysis are available at https://github.com/Sorennorge/Choroid-plexus-orthology-study. The sequencing data for human and mouse were quality controlled with fastqc (Andrews, 2010), and the human data were trimmed with Trimmomatic (Bolger *et al*, 2014) (Sliding window 4:20, minimum length of 35 bp). The obtained sequencing data were mapped to reference genome for human (Homo sapiens GRCh38 v.104), rat (Rattus norvegicus Rnor_6.0 v.104), mouse (Mus musculus GRCm39 v.104), and zebrafish (Danio rerio GRCz11 v.104) using Spliced Transcripts Alignment to a Reference (STAR) RNA-seq aligner (v. 2.7.2a) (Dobin *et al*, 2013). The reference genomes were trimmed prior to mapping to only contain gene/transcripts of biotype ‘protein coding’. The mapped alignment by STAR was normalized to TPM with RSEM (RNA-Seq by Expectation Maximization v. 1.3.3) (Li & Dewey, 2011). The means of calculated TPM were used for the further analysis of human, mouse, and zebrafish. Especially, the TPMs of zebrafish presented in SuppleTable 2 are averaged counts from the 2 sequencing datasets. Gene information for ‘gene symbol’ (gene names) and orthologs were gathered from Ensembl biomart (Cunningham *et al*, 2022; Kinsella *et al*, 2011). The density analysis was generated utilizing the seaborn distribution python library (Hunter, 2007; Waskom, 2021). The Venn diagrams were generated utilizing nVennR package for R (Perez-Silva *et al*, 2018). Transporters were collected based on the protein class which was obtained by utilizing the Panther database (Mi *et al*, 2021) with gene symbols. Transporters were ranked according to their TPM expression within each species. Hereafter, the transporter table were sorted based on the mean rank for all four species.

### Cloning zebrafish *fabp7b* and whole mount RNA *in situ* hybridization

To isolate *fabp7b* gene, primers were designed to clone *fabp7b* ORF from Ensembl (*fabp7b*: ENSDARG00000034650). *Fabp7b* ORF was amplified from 3 dpf cDNA using the designed primers by PCR and cloned into pGEM^®^-T easy vectors (Promega). To synthesize fabp7b RNA probe, the cloned vector was linearized by SacII restriction enzymes (New England Biolabs) at 37 °C for 1 hour in 30 μL solution (3 μL CutSmart buffer, 2 μL SacII enzyme and 25 μL fapb7b ORF cloned vectors) and then purified by the ethanol precipitation described in the section of total RNA preparation above. The linearized purified vectors were transcribed using the SP6 RNA polymerase and DIG-oxygenin labeling mixture (Roche) for 1 hour at 37°C in 20 μL mixture (2 μL 10x transcription buffer, 2 μL 10x DIG RNA labelling mix, 1 μL RNase inhibitor, 2 μL SP6 polymerase, and 13 μL linearized plasmid. The synthesized RNA probe was treated with 2 μL Dnase for 10 min at 37°C to remove left DNA plasmid. Then the probe was purified by ethanol precipitation and dissolved in 20 μL formamide and stored at −20°C. The RNA probe integrity was confirmed by gel electrophoresis and the probe quantity was measured by NanoDrop (Thermo Scientific).

To prepare samples for whole mount RNA *in situ* hybridization, 7 dpf zebrafish larvae were euthanized by MS222 (Sigma-Aldrich) in E3 medium until movements ended and fixed with 4% paraformaldehyde (PFA) in 0.2 % PBST (PBS with 0.2% Triton-X100) overnight at 4 degrees. The fixed larvae were rinsed with PBST several times and the brains were dissected out from the larvae with fine forceps. The larval brains were dehydrated in 100% methanol (Merk) and kept at −20°C.

RNA *in situ* hybridization was performed by alkaline phosphatase-NBT/BCIP (Roche) chromogenic reaction, as described earlier (Jeong *et al*., 2021; Thisse & Thisse, 2008). The brains were rehydrated gradually through 75%, 50%, 25% methanol in 0.2 % PBST solution steps for 5 min per each solution. Following 3 washes for 15 min in 0.2 % PBST, the samples were digested with proteinase K (10 μg/mL) for 10-20 secs, fixed in 4% PFA solution for 20 min at RT and then washed by 0.2 % PBST. The samples were prehybridized at 65°C for 3 hours in hybridization buffer (50% formamide (Junsei), 5X SSC (Sigma-Aldrich), 50 μg/mL heparin, 500 μg/mL tRNA, 0.2% Triton X-100), followed by hybridization at 60°C overnight with 200 ng RNA probe / 200 μL hybridization buffer. The samples were subsequently washed twice for 20 min in 50% formamide/2X SSCT, twice for 15 min in 2X SSCT, three times for 20 min in 0.2X SSCT and 0.1XSSCT once for 10 min at 65°C. Following three washes in 0.1% PBT (0.1% tween-20 in PBS) at RT on shaker, the samples were blocked in blocking solution (5% sheep serum, 2 mg/mL BSA) for 2 hours at RT on shaker and incubated overnight at 4°C in AP-anti-DIG antibody (Roche) diluted 1:4000 in blocking solution. The next day, the samples were washed 10 x15 min in 0.1% PBT, followed by 3 x 5 min in staining buffer (100 mM Tris, pH 9, 50 mM MgCl2, 100mM NaCl, 0.1% Tween-20) at RT. The samples were stained for 4-12 hours in NBT/BCIP solution (1:50 dilution of NBT/BCIP (Roche) in staining buffer). The staining was terminated by several washes with 0.1% PBT at RT and fixation with 4% PFA solution overnight at 4°C. The samples were washed with 0.1% PBT several times at RT and transferred to 75% glycerol in PBS. The brains were imaged in the bright field mode of Ni-U fluorescent microscope (Nikon) with DS-Ri2 camera.

### Metronidazole treatment

To ablate the epithelial cells in the ChP, juvenile *Gt(fabp7b:2A-Gal4vp16);Tg(uas:NTR-mCherry);Gt(foxj1b:GFP)* zebrafish and their mCherry negative *Gt(foxj1b:GFP)* siblings were used. 30 larvae from each line were kept in a nursery tank (Techniplast) before metronidazole (MTZ) treatment to produce similar size of the juvenile fish. At 14 dpf, *Gt(fabp7b:2A-Gal4vp16);Tg(uas:NTR-mCherry);Gt(foxj1b:GFP)* and mCherry negative *Gt(foxj1b:GFP)* fish were treated with either 10 mM MTZ (Sigma-Aldrich) in 0.2% DMSO in AFW for 24 hour (from 14 to 15 dpf). As another control group, *Gt(fabp7b:2A-Gal4vp16);Tg(uas:NTR-mCherry);Gt(foxj1b:GFP)* fish were also treated with 0.2% DMSO in AFW for 24 hours (Agetsuma *et al*., 2010; Palumbo *et al*, 2020). Three fish were placed in a petri dish containing 50 mL of 10 mM MTZ / 0.2% DMSO in AFW or 0.2% DMSO in AFW in a 28°C incubator, with regular light/dark cycles (14 hours light / 10 hours dark). Following treatment, the fish were washed with fresh AFW three times and transferred to the nursery tank until the start of the experiments. MTZ treatment experiments were performed on three different batch of fish on different dates. Data were collected and analyzed from the three experiments.

### Intravenous injection and wholemount zebrafish *in vivo* live imaging

For intravenous injection, juvenile fish (17-18 dpf) were anesthetized in MS222 diluted in AFW and mounted in 1.5 % low melting agarose with the trunk veins facing upwards. The body size of each fish was measured with a ruler. 10 kDa dextran Alexa Fluor 647 (Thermo Fisher Scientific) was dissolved in PBS at a final concentration of 10 mg/mL (O’Brown *et al*, 2019). Intravenous injections were performed as described above for the ventricular injections: injection needles were pulled with the identical method and the needle tip was opened by cutting with forceps. A microinjector (Eppendorf Femtojet 4i) was used to microinject circa 2-3 nL of the fluorescent dye into the cardinal vein near the tail. Following injection, the fish were released from agarose and placed in MS222 diluted AFW for 1 hour. Then, the fish were remounted in 1.5 % low melting agarose with the dorsal brain facing upwards to image the tel- or rhombencephalon areas including forebrain and hindbrain ChP. The fish were transferred to the confocal microscope and Z-stacks images were obtained with 2.5 μm z-steps by a 20x water-immersion objective (Zeiss, NA 1.0, Plan-Apochromat) at RT.

### Imaging processing

Confocal stacks were stitched in Fiji/ImageJ using the pairwise stitching plugin (Preibisch *et al*, 2009) for images shown in Figure 3C1, Supplementary Figure 5B, 6A1. All fluorescence images except the stitched images were deconvoluted in the Huygen software (Scientific Volume Imaging) using the standard Deconvolution Express function. Raw images were used for all analysis and quantifications.

### Quantification and statistical analysis

All quantifications were performed using by ImageJ/Fiji. Plots and statistical analyses were generated using MATLAB. The statistical tests used and description of the numbers of observations are indicated in each figure legend.

#### Quantification of NTR-mCherry expression in the tela choroidea and choroid plexus

To measure the percentage of NTR-mCherry-expressing areas in each choroid plexus, the confocal stacks from the DMSO treated control *Gt(fabp7b:2A-Gal4vp16);Tg(uas:NTR-mCherry);Gt(foxj1b:GFP)* fish at 17-18 dpf (*n* = 10) were utilized. First, max projection images of the confocal stacks were generated by ImageJ/Fiji which contained GFP-expressing the tela choroidea, forebrain and hindbrain ChPs. Then, the region of interest (ROI) in GFP and mCherry-expressing area was drawn manually for each fish and recorded. The size of the mCherry-expressing area was normalized to the GFP-expressing area for each fish. The values were converted to percentages. All datapoints were imported and plotted in MATLAB.

To quantify the ablation efficiency in the tela choroidea, forebrain and hindbrain ChPs, max projection images of the confocal stacks were generated from control and ChP ablation groups. A ROI comprising the GFP-positive area was drawn for each fish. The GFP-positive area was normalized by the average value of all GFP-positive area obtained from controls (DMSO and Ctrl MTZ treated groups). All values were imported and plotted in MATLAB.

#### Quantification of the size of brain ventricles on confocal images

We quantified the size of the anterior telencephalic ventricle (ant. TV) on images showing a dorsal view, and the posterior telencephalic/anterior di-/mesencephalic ventricle (post. TV & D/MV) and rhombencephalic ventricle (RV) on resliced images showing the sagittal view. We selected these view angles as they allowed us to distinguish the brain ventricles from blood plasma and skin.

For measuring the ant. TV areas, 12.5 µm thick (6 slices) maximum intensity projections of confocal stacks were used. For the rest of the ventricular areas, images of the confocal stacks were resliced by ImageJ/Fiji to visualize the sagittal view of the brain. Then 10 µm thick (5 slices) maximum intensity projection of the stacks were utilized. ROIs were drawn on the far-red channel showing Alexa 647 positive ventricles. The values were imported and plotted in MATLAB.

#### Quantification of the fluorescent intensities on confocal images

To quantify the fluorescent intensities in the blood, ventricle, and parenchyma, we used the same sections and areas of the brain where the ventricular size was measured. Mean fluorescent intensities were measured by ImageJ from ROIs drawn on blood vessels, parenchyma and ventricle for each section and each area. As a result, for each fish, 6 or 5 fluorescent intensities were collected per area. Intensities were normalized to the size of the ROI and averaged to obtain the “mean intensities per area”. The values in the parenchyma (Intensity_parenchyma_) and ventricle (Intensity_ventricle_) were further normalized by the values in the blood (Intensity_blood_). All datapoints were collected and imported to MATLAB.

#### Quantification of the brain parenchyma and fish size on confocal images

To check the size of brain parenchyma in the control and ChP ablated groups, the width of telencephalon and rhombencephalon was measured by ImageJ/Fiji. The tel-/rhombencephalic width was the widest area observed at the tel-/rhombencephalon. The rhombencephalon, parenchyma width was measured where the RV was widest. The values were imported and plotted in MATLAB. The fish size was measured using a ruler before intravenous injection and these datapoints were also imported and plotted to MATLAB.

#### Figure assembly

All figures were assembled using Adobe Illustrator.

