## Supplemental Figures for "The evolutionary conserved choroid plexus sustains the homeostasis of brain ventricles in zebrafish"

### Supplemental information

### Supplementary Figure legends

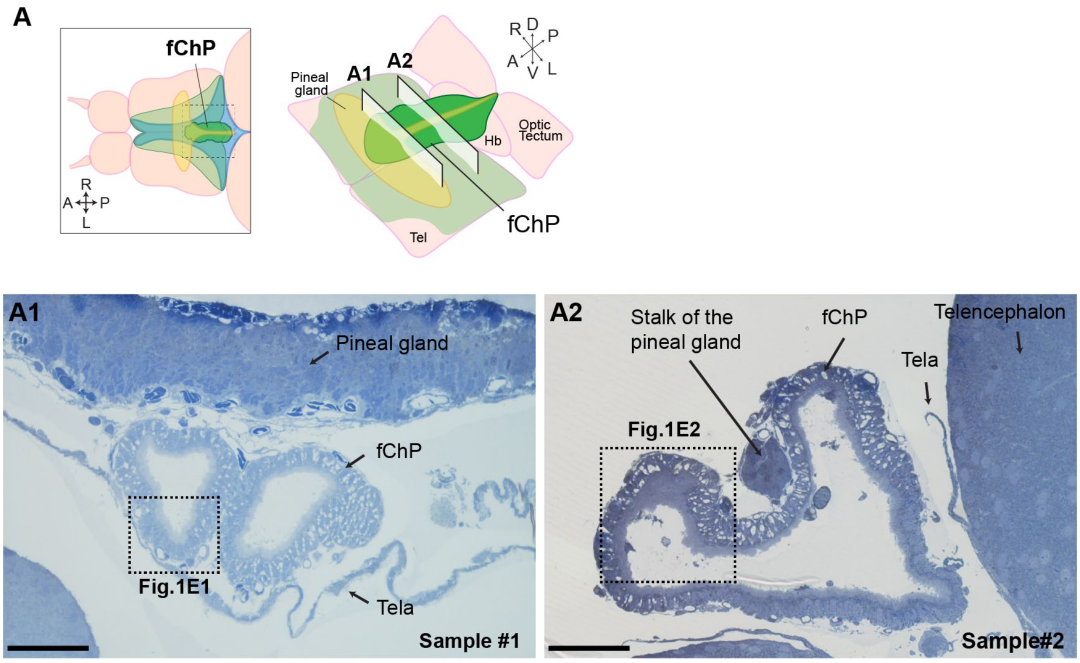

**Supplementary Figure 1. Cross section views of the adult forebrain choroid plexus used for electron microscopy. Related to Figure 1.**

**(A)** Schematic of the forebrain choroid plexus (fChP) in the adult brain explant for transmission electron microscopy. **(A1-A2)** Light transmission images of the cross sections shown in Fig.1E1-1E2. The sample#1 and #2 are used in Fig.1E1 and Fig.1E2, respectively. (*n* = 2)

Abbreviations: A, anterior; P, posterior; D, dorsal; V, ventral; R, right; L, left; Hb, habenula; Tel, telencephalon. Scale bars: 50 μm.

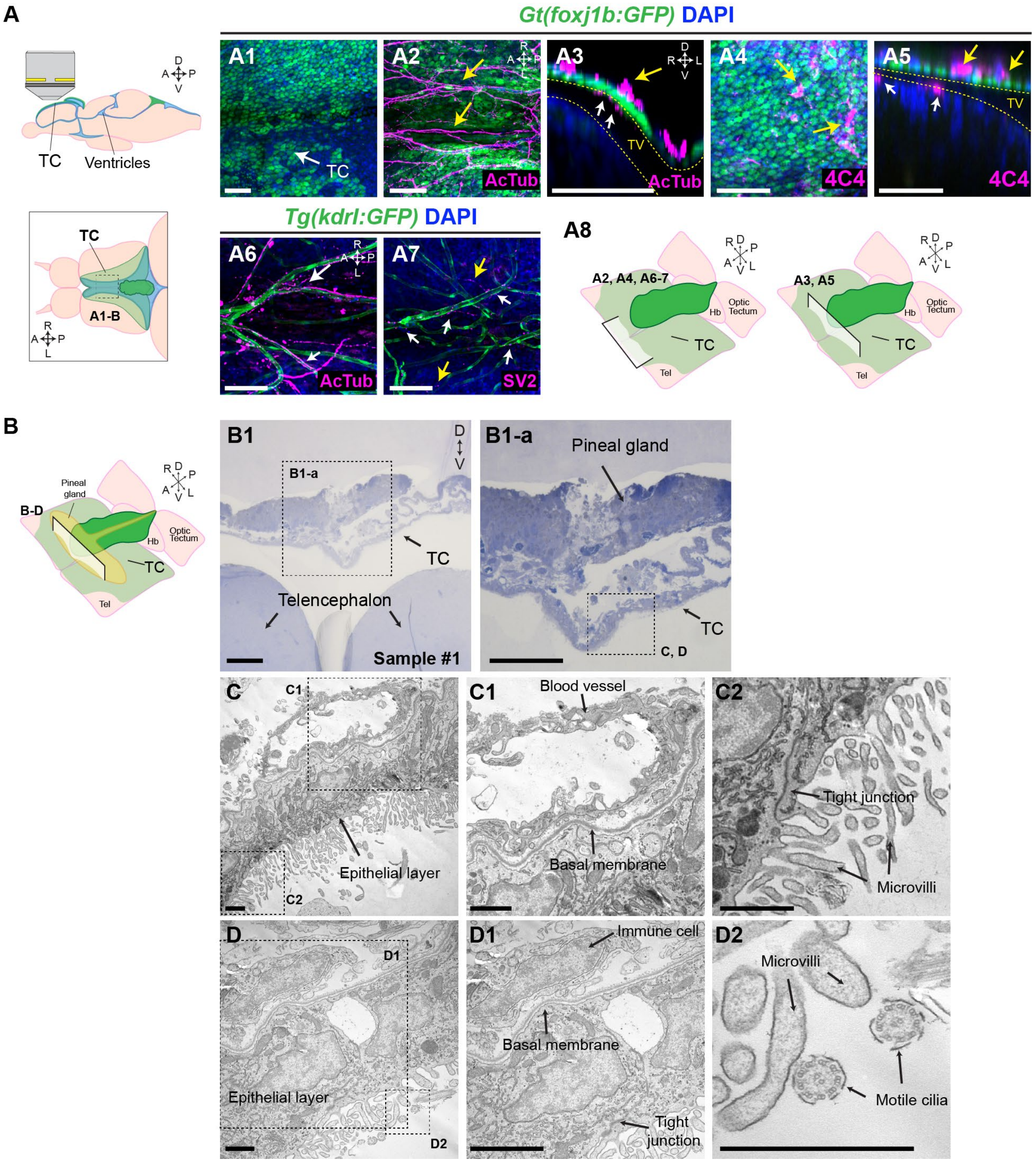

**Supplementary Figure 2. The tela choroidea has similar tissue characteristics as the choroid plexus. Related to Figure 1.**

**(A)** Diagram of the adult brain explant for imaging the tela choroidea (TC). Lateral (upper) and dorsal (lower) views of the brain. **(A1-A5)** The *Gt(foxj1b:GFP)* labels epithelial cells of the TC. DAPI labels nuclei. **(A1)** Max projection of the TC. **(A2-A3)** Max projection **(A2)** and single plane **(A3)** images of the TC immunolabelled by acetylated tubulin (AcTub) antibody labelling both cilia (white arrows) and axons (yellow arrows). **(A4-A5)** 4C4 antibody labelling of immune cells in the TC. **(A6-A7)** Max projection (dorsal view) images of the TC in the *Tg(kdrl:GFP)* immunolabelled by GFP, AcTub **(A6)** and SV2 antibodies for presynaptic vesicles **(A7)** with DAPI. **(A8)** Schemes representing the location of the images shown in (A2-A7).

**(B)** Diagram of the TC in the adult brain explant using for transmission electron microscopy.

**(B1, B1-a)** Low and high magnification light transmission images of the cross sections using in panel C-D2.

**(C-D)** Transmission electron microscopy of the TC. High magnification images **(C1-C2, D1-D2)** from dotted boxes in (C, D).

Abbreviations: A, anterior; P, posterior; D, dorsal; V, ventral; R, right; L, left. Scale bars: (A1-B1-a), 50 µm; (C-D2), 1 µm.

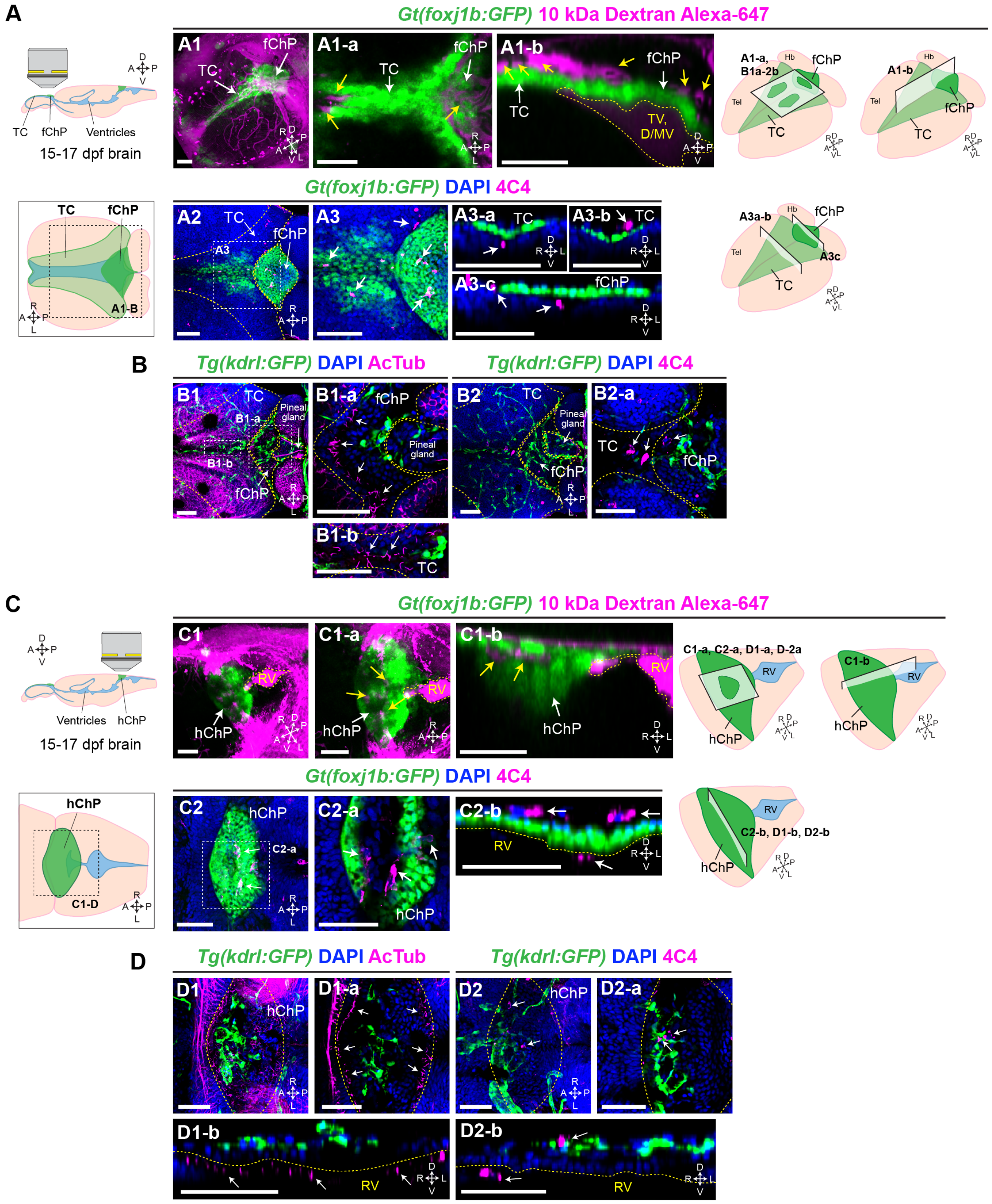

**Supplementary Figure 3. The 2 weeks-old juvenile choroid plexus is less mature than in the adult stage.**

**(A)** Diagram of the juvenile brain for imaging the TC and fChP at 15-17 days post fertilization (dpf). Lateral (upper) and dorsal (lower) views of the brain. **(A1-A1-b)** Images of the tel-/diencephalon in the *Gt(foxj1b:GFP)* injected with a 10 kDa Alexa-647 fluorescent dye at 17 dpf to visualize the blood vessels. Left: scheme representing the location of individual panels shown in (A1-B2). **(A1)** 3D reconstructed image and **(A1-a, A1-b)** single plane images of the TC-fChP. Yellow arrows indicate blood vessels nearby the TC or fChP (white arrows). **(A2-A3-c)** *Gt(foxj1b:GFP)* juvenile brain immunolabelled by 4C4 and GFP antibodies with DAPI. White arrows label 4C4+ immune cells in the TC-fChP. **(A2-A3)** Max projection images of the TC-fChP. **(A3-a-A3-c)** Single plane images of the TC-fChP.

**(B)** *Tg(kdrl:GFP)* juvenile brain immunolabelled by acetylated tubulin (AcTub) or 4C4 antibodies with GFP antibody. **(B1, B2)** Max projection images of the TC-fChP. **(B1-a, B1-b)** Max projection images of confocal stacks of the fChP **(B1-a)** and TC **(B1-b)**. White arrows indicate cilia. **(B2-a)** Max projection images of confocal stacks of the TC-fChP. White arrows indicate 4C4+ immune cells.

**(C)** Diagram of the juvenile brain for imaging the hChP at 17 dpf. Lateral (upper) and dorsal (lower) views of the brain. **(C1-C1-b)** Images of the rhombencephalon in the *Gt(foxj1b:GFP)* injected with 10 kDa Alexa-647 fluorescent dye at 17 dpf. Left: scheme representing the location of individual panels shown in (C1-D2). **(C1)** 3D reconstructed image of confocal stacks and **(C1-a, C1-b)** single plane images of the hChP (C1-a, C1-b). Yellow arrows indicate blood vessels nearby the hChP (white arrows). **(C2-C2-b)** *Gt(foxj1b:GFP)* juvenile brain explants immunolabelled by 4C4 and GFP antibodies. White arrows label 4C4+ immune cells in the hChP. **(C2)** Max projection of the hChP and **(C2-a-C2-b)** single plane images

**(D)** *Tg(kdrl:GFP)* juvenile brain explants immunolabelled by AcTub or 4C4 antibodies with GFP antibody. White arrows indicate 4C4+ immune cells. White arrows indicate cilia in (D1-D1-b) and 4C4+ immune cells in (D2-D2-b).

Yellow dotted lines mark the tissue boundaries such as TC, fChP or ventricles. DAPI labels nuclei.

Abbreviations: TC, tela choroidea; fChP, forebrain choroid plexus; hChP, hindbrain choroid plexus, TV, telencephalic ventricle; D/MV, di-/mesencephalic ventricle; RV, rhombencephalic ventricle; A, anterior; P, posterior; D, dorsal; V, ventral; R, right; L, left. Scale bars: 50 μm.

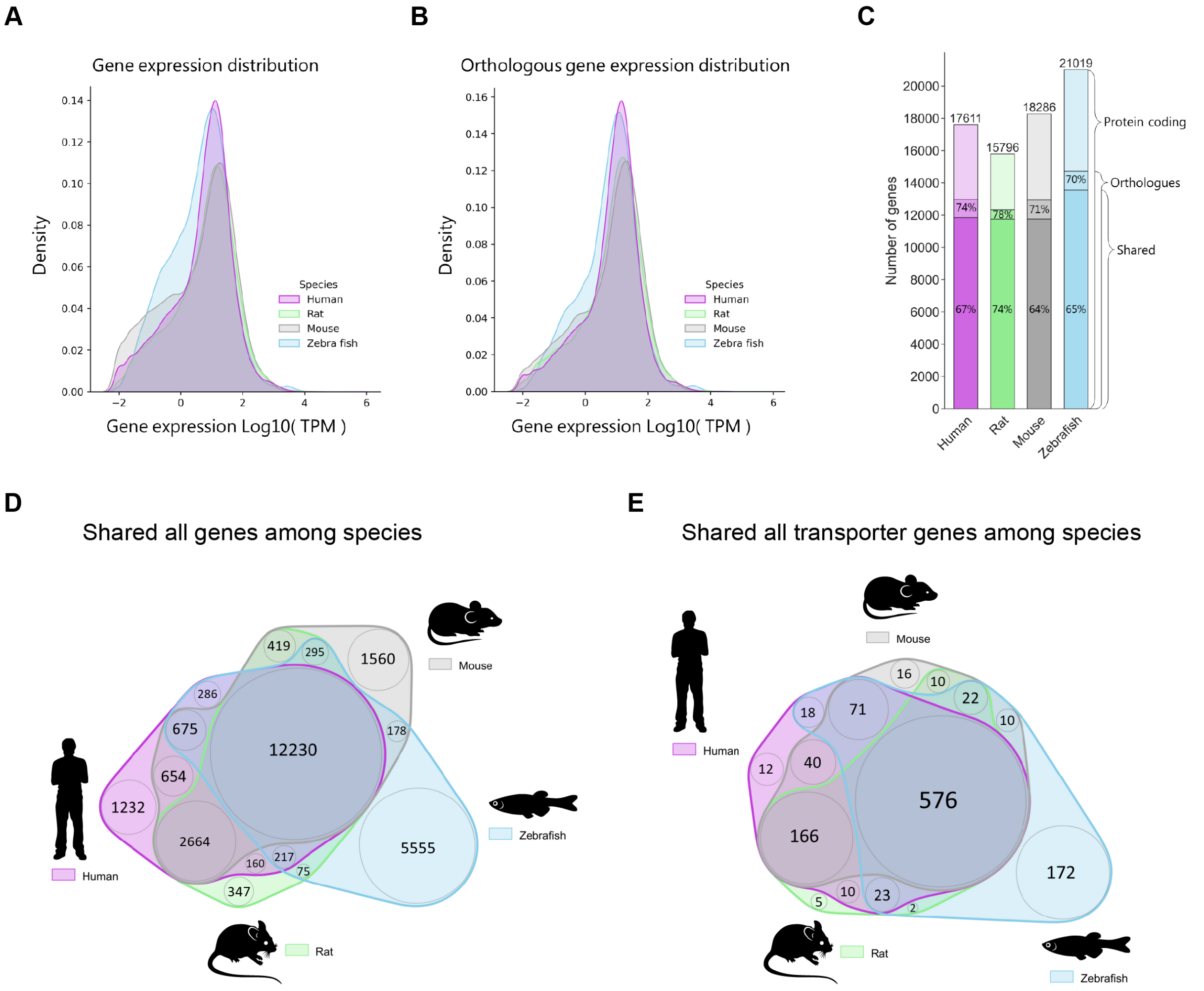

**Supplementary Figure 4. Gene expression of the choroid plexus across vertebrates. Related to Figure 2.**
**(A)** Distribution of the gene expression levels for all protein coding genes for the four species.
**(B)** Distribution of the gene expression levels for the protein coding genes that have an ortholog among the four species.
**(C)** The protein coding gene count for human, rat, mouse, and zebrafish, including the percentage of protein coding genes that have orthologs in any of the other species and the percentage of protein coding genes that have an ortholog in all four species.
**(D)** Weighted Venn diagram showing protein coding genes expressed in the ChP of the four species.
**(E)** Weighted Venn diagram showing transporter genes shared expressed in the ChP of the four species.
Abbreviations: TPM, transcript per million

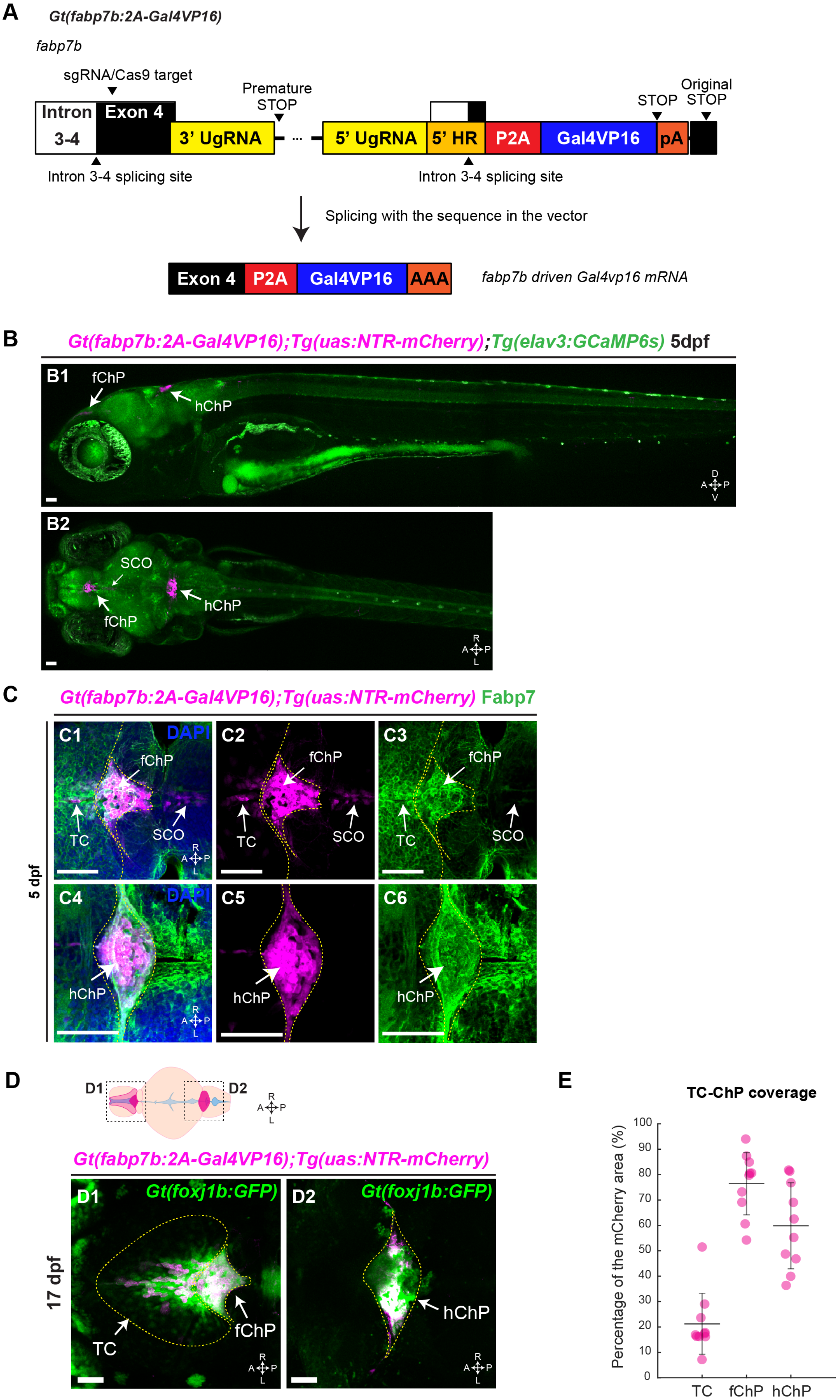
A
Gt(fabp7b:2A-Gal4VP16)
fabp7b
sgRNA/Cas9 target
Premature STOP
Original STOP
STOP
Intron 3-4
Exon 4
3' UgRNA
5' UgRNA
5' HR
P2A
Gal4VP16
pA
Intron 3-4 splicing site
Intron 3-4 splicing site
Splicing with the sequence in the vector
Exon 4
P2A
Gal4VP16
AAA
fabp7b driven Gal4vp16 mRNA
B
Gt(fabp7b:2A-Gal4VP16);Tg(uas:NTR-mCherry);Tg(elav3:GCaMP6s) 5dpf
B1
fChP
hChP
B2
SCO
fChP
hChP
C
Gt(fabp7b:2A-Gal4VP16);Tg(uas:NTR-mCherry) Fabp7
5 dpf
C1
DAPI
C2
C3
C4
C5
C6
fChP
TC
SCO
hChP
D
D1
D2
Gt(fabp7b:2A-Gal4VP16);Tg(uas:NTR-mCherry)
17 dpf
Gt(foxj1b:GFP)
TC
fChP
hChP
E
TC-ChP coverage
Percentage of the mCherry area (%)
TC
fChP
hChP

**Supplementary Figure 5. Generation of the *fabp7b* specific knockin zebrafish. Related to Figure 3.**

**(A)** Schematic of *fabp7b* gene locus in the *Gt(fabp7b:2A-Gal4vp16)* line. In this line, the targeting vector plasmid was integrated into the genome, which produces a premature stop codon upstream of the *P2A-Gal4vp16* coding sequence. The splicing site in the homology arm of the target vector allows to still generate *fabp7b-P2A-Gal4vp16* mRNA, via alternative splicing.

**(B-B2)** Confocal images of an entire 5 dpf *Gt(fabp7b:2A-Gal4vp16);Tg(uas:NTR-mCherry);Tg(elavl3:GCaMP6s)* larvae. Lateral **(B1)** and dorsal **(B2)** view of the line.

**(C)** Dorsal view of a 5dpf *Gt(fabp7b:2A-Gal4vp16);Tg(uas:NTR-mCherry)* larval brain immunolabelled with Fabp7/BLBP antibody.

**(D)** Diagram of the juvenile brain used for imaging the TC-ChPs as shown in (D1-D2). **(D1-D2)** Maximum projection of images of the *Gt(fabp7b:2A-Gal4vp16);Tg(uas:NTR-mCherry);Gt(foxj1b:GFP)* at 17 dpf.

**(E)** Quantification of the coverage rate of the TC, fChP and hChP areas by the *Gt(fabp7b:2A-Gal4vp16);Tg(uas:NTR-mCherry)*. The TC, fChP and hChP areas were defined as the foxj1b-GFP positive areas. $n = 10$.

Abbreviations: TC, tela choroidea; fChP, forebrain choroid plexus; hChP, hindbrain choroid plexus; A, anterior; P, posterior; D, dorsal; V, ventral; R, right; L, left. Scale bars: 50 μm.

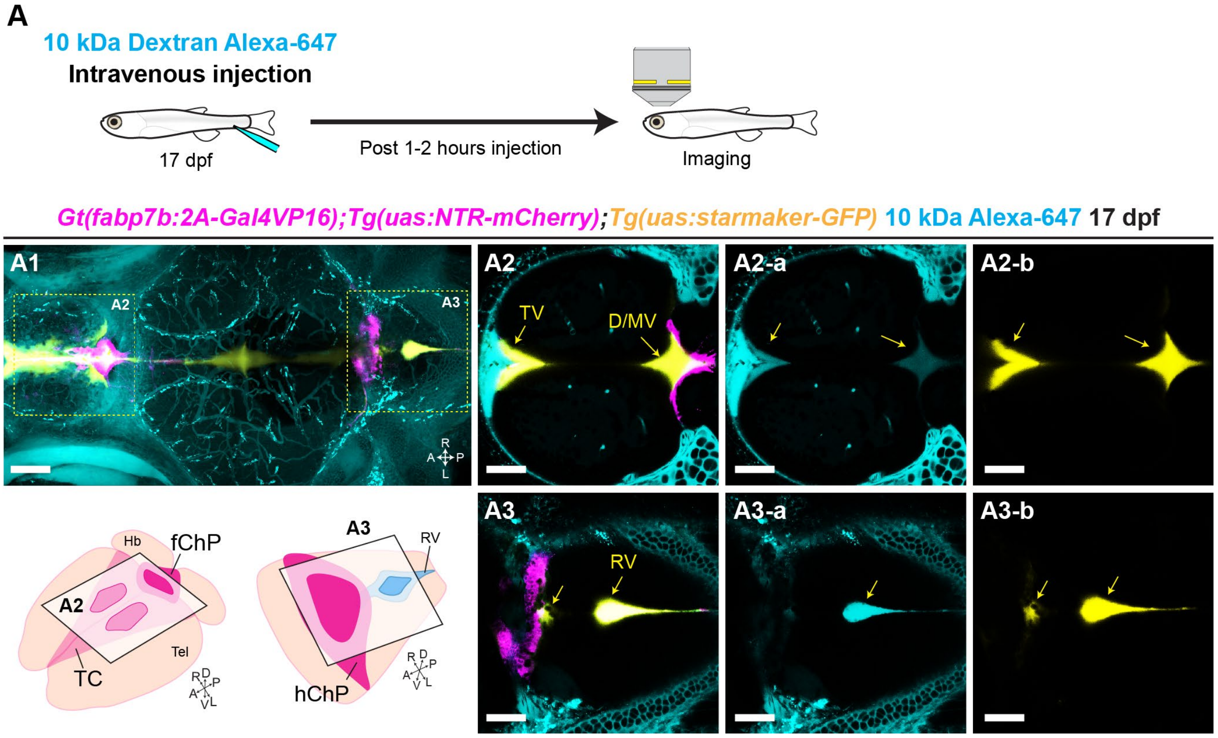

**Supplementary Figure 6. Intravenous injection of fluorescent dyes labels both the blood plasma and the brain ventricles. Related to Figure 4.**

**(A)** Experimental scheme used to visualize both the blood plasma and the brain ventricles after injecting 10 kDa Alexa-647 fluorescent dye into the cardinal vein of the trunk at 17dpf. **(A1)** Maximum projection of the *Gt(fabp7b:2A-Gal4vp16);Tg(uas:NTR-mCherry);Tg(uas:starmaker-GFP)* line injected 10 kDa Alexa-647 fluorescent dye at 17 dpf. Yellow dotted boxes show single plane of each area, telencephalon (A2) and rhombencephalon (A3). In the lower panel are shown schemes representing the location of the images shown in (A2-A3). **(A2, A3)** Merged images of Gal4vp16 driven NTR-mCherry (magenta) and starmaker-GFP (yellow) and intravenously injected the Alexa-647 fluorescent dye (cyan). Yellow arrows mark each brain ventricle, labelled by both starmaker-GFP and Alexa-647 dye. **(A2-a, A2-b, A3-a, A3-b)** Single channel images of intravenously injected Alexa-647 fluorescent dye **(A2-a, A-3a)** and starmaker-GFP **(A2-b, A3-b)**.

Abbreviations: TC, tela choroidea; fChP, forebrain choroid plexus; hChP, hindbrain choroid plexus; TV, telencephalic ventricle; D/MV, dien-/mesencephalic ventricle; RV, rhombencephalic ventricle; A, anterior; P, posterior; D, dorsal; V, ventral; R, right; L, left. Scale bars: 50 µm.

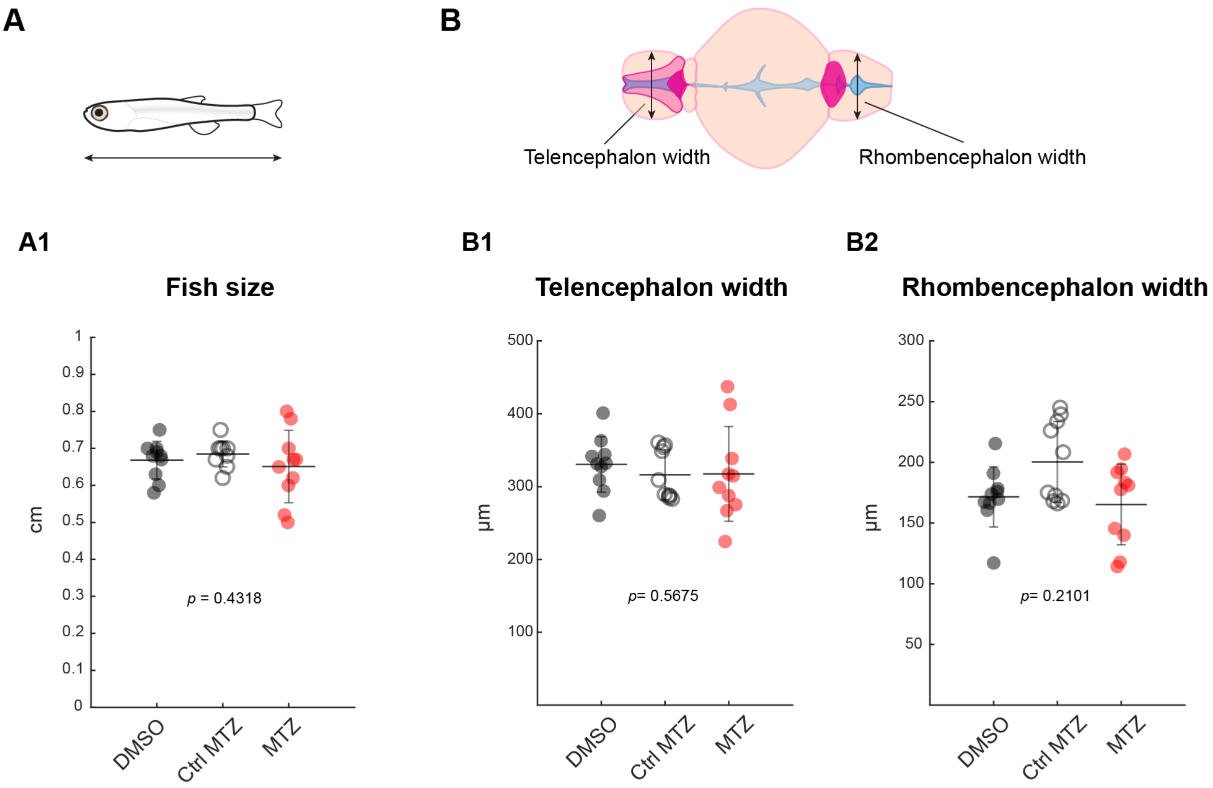

**Supplementary Figure 7. Brain and body size are not affected in the choroid plexus ablated zebrafish. Related to Figure 4.**

**(A)** Diagram of the 2 weeks-old juvenile zebrafish. The arrow bar indicates the length of the fish measured by a ruler. **(A1)** Quantification of the fish size between control (DMSO, Ctrl MTZ) and TC-ChP-ablated fish (MTZ).

**(B)** Diagram of the 2 weeks-old juvenile brain (dorsal view). The arrow bars indicate the width of the telencephalon and rhombencephalon parenchyma. **(B1-B2)** Quantification of the brain parenchyma width between control and TC-ChP-ablated fish.

Statistical analysis: Each group; $n$ =10, Kruskal-Wallis test, $p > 0.05$.

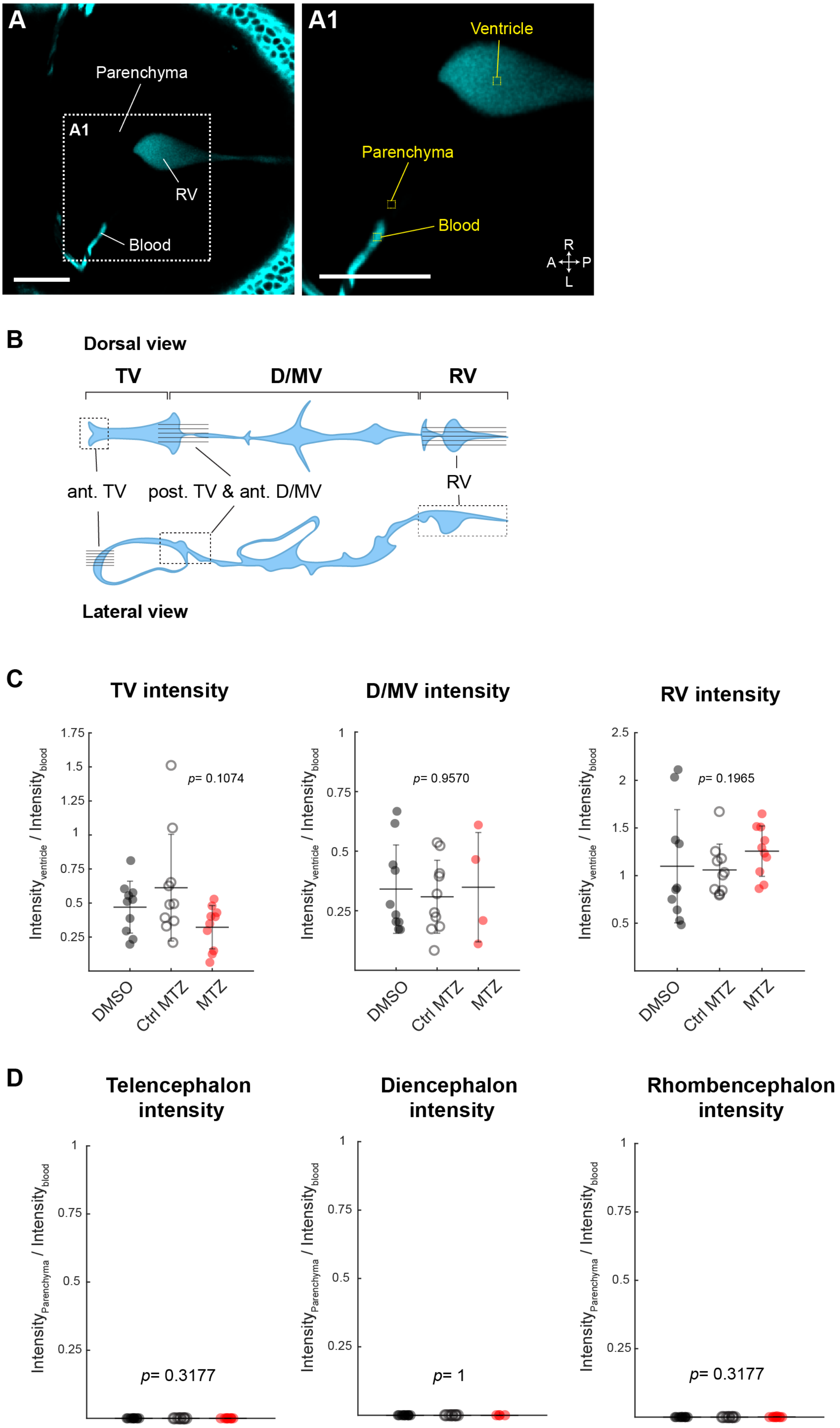

A
Parenchyma
A1
RV
Blood
A1
Ventricle
Parenchyma
Blood
R
A
P
L
B
Dorsal view
TV
D/MV
RV
ant. TV
post. TV & ant. D/MV
RV
Lateral view
C
TV intensity
D/MV intensity
RV intensity
p= 0.1074
p= 0.9570
p= 0.1965
DMSO
Ctrl MTZ
MTZ
D
Telencephalon intensity
Diencephalon intensity
Rhombencephalon intensity
p= 0.3177
p= 1
p= 0.3177

**Supplementary Figure 8. No leakage of blood tracers in the CSF and brain parenchyma is observed in the choroid plexus ablated zebrafish. Related to Figure 4.**

**(A-A1)** Representative image showing how fluorescent intensities of blood plasma, brain parenchyma and ventricle were measured. Single plane of the rhombencephalon area (dorsal view) in control fish is shown in (A1).

**(B)** Diagram of the brain ventricles in the juvenile brain. Dotted boxes and back lines are the confocal stacks where fluorescent intensities were measured.

**(C-D)** Quantification of fluorescent intensities in the brain ventricle **(C)** or brain parenchyma **(D)** between control (DMSO, Ctrl MTZ) and TC-ChP-ablated fish (MTZ).

Statistical analysis: Each control group, n = 10; TC-ChPs ablated group $n$ = 4 in the D/MV and diencephalon intensity due to collapse of the ventricle. Kruskal-Wallis test, $p > 0.05$. Abbreviations: TV: telencephalic ventricle; D/MV: dien-/mesencephalic ventricle; RV: rhombencephalic ventricle; ant.: anterior; post.: posterior. Abbreviations: A, anterior; P, posterior; D, dorsal; V, ventral; R, right; L, left. Scale bars: 50 μm

**Supplementary Table 1. Sample preparation protocol using Pelco Biowave Pro+ for electron microscopy. Related to Figure 1.**

| STEP | Description | TIME *H:MIN:SEC* | USER PROMPT | Power *Watts* | STEADY TEMP °C | MAX TEMP° C *(limit temp)* | VACUUM CYCLE *Auto=ON (Trykk 20, vacuum time 30, vent time 30)* | Solution (concentration) |
|---|---|---|---|---|---|---|---|---|
| 1 | Rinse in buffer | 00:00:40 | 0FF | 250 | 20 | 50 | OFF | |
| 2 | Rinse in buffer | 00:00:40 | OFF | 250 | 20 | 50 | OFF | 2 % in 0,15 M cacodylate buffer (pH 7.4) |
| 3 | Osmium ON | 00:02:00 | ON | 100 | 20 | 50 | AUTO | |
| 4 | Osmium OFF | 00:02:00 | OFF | 0 | 20 | 50 | AUTO | |
| 5 | Osmium ON | 00:02:00 | OFF | 100 | 20 | 50 | AUTO | |
| 6 | Osmium OFF | 00:02:00 | OFF | 0 | 20 | 50 | AUTO | |
| 7 | Osmium ON | 00:02:00 | OFF | 100 | 20 | 50 | AUTO | |
| 8 | Osmium OFF | 00:02:00 | OFF | 0 | 20 | 50 | AUTO | |
| 9 | Osmium ON | 00:02:00 | OFF | 100 | 20 | 50 | AUTO | |
| 10 | KFC ON | 00:02:00 | ON | 100 | 20 | 50 | AUTO | 2,5 % in 0,15 M cacodylate buffer (pH 7.4) |
| 11 | KFC OFF | 00:02:00 | OFF | 0 | 20 | 50 | AUTO | |
| 12 | KFC ON | 00:02:00 | ON | 100 | 20 | 50 | AUTO | |
| 13 | KFC OFF | 00:02:00 | OFF | 0 | 20 | 50 | AUTO | |
| 14 | KFC ON | 00:02:00 | ON | 100 | 20 | 50 | AUTO | |
| 15 | KFC OFF | 00:02:00 | OFF | 0 | 20 | 50 | AUTO | |
| 16 | KFC ON | 00:02:00 | ON | 100 | 20 | 50 | AUTO | |
| 17 | Rinse in water Fume hood, 2 times | | ON | 0 | 20 | 50 | OFF | |
| 18 | Rinse in water | 00:00:40 | ON | 250 | 20 | 50 | OFF | |
| 19 | Rinse in water | 00:00:40 | ON | 250 | 30 | 50 | OFF | |
| 20 | Pyrogallol ON | 00:02:00 | ON | 100 | 40 | 50 | AUTO | 320 mM in aqueous solution (pH 4.1), filtered |
| 21 | Pyrogallol OFF | 00:02:00 | OFF | 0 | 40 | 50 | AUTO | |
| 22 | Pyrogallol ON | 00:02:00 | OFF | 100 | 40 | 50 | AUTO | |
| 23 | Pyrogallol OFF | 00:02:00 | OFF | 0 | 40 | 50 | AUTO | |
| 24 | Pyrogallol ON | 00:02:00 | OFF | 100 | 40 | 50 | AUTO | |
| 25 | Pyrogallol OFF | 00:02:00 | OFF | 0 | 40 | 50 | AUTO | |
| 26 | Pyrogallol ON | 00:02:00 | OFF | 100 | 40 | 50 | AUTO | |
| 27 | Rinse in water Fume hood, 2 times | | ON | 0 | 20 | 50 | OFF | |
| 28 | Rinse in water | 00:00:40 | ON | 250 | 20 | 50 | OFF | |
| 29 | Rinse in water | 00:00:40 | ON | 250 | 20 | 50 | OFF | |
| 30 | Osmium ON | 00:02:00 | ON | 100 | 20 | 50 | AUTO | 1% OsO4 in aqueous solution |
| 31 | Osmium OFF | 00:02:00 | OFF | 0 | 20 | 50 | AUTO | |
| 32 | Osmium ON | 00:02:00 | OFF | 100 | 20 | 50 | AUTO | |
| 33 | Osmium OFF | 00:02:00 | OFF | 0 | 20 | 50 | AUTO | |
| 34 | Osmium ON | 00:02:00 | OFF | 100 | 20 | 50 | AUTO | |
| 35 | Osmium OFF | 00:02:00 | OFF | 0 | 20 | 50 | AUTO | |
| 36 | Osmium ON | 00:02:00 | OFF | 100 | 20 | 50 | AUTO | |
| 37 | Rinse in water Fume hood, 2 times | | ON | 0 | 20 | 50 | OFF | |
| 38 | Rinse in water | 00:00:40 | ON | 250 | 20 | 50 | OFF | |
| 39 | Rinse in water | 00:00:40 | ON | 250 | 30 | 50 | OFF | |
| 40 | Pyrogallol ON | 00:02:00 | ON | 100 | 40 | 50 | AUTO | 320 mM in aqueous solution (pH 4.1), filtered |
| 41 | Pyrogallol OFF | 00:02:00 | OFF | 0 | 40 | 50 | AUTO | |
| 42 | Pyrogallol ON | 00:02:00 | OFF | 100 | 40 | 50 | AUTO | |
| 43 | Pyrogallol OFF | 00:02:00 | OFF | 0 | 40 | 50 | AUTO | |
| 44 | Pyrogallol ON | 00:02:00 | OFF | 100 | 40 | 50 | AUTO | |
| 45 | Pyrogallol OFF | 00:02:00 | OFF | 0 | 40 | 50 | AUTO | |
| 46 | Pyrogallol ON | 00:02:00 | OFF | 100 | 40 | 50 | AUTO | |
| 47 | Rinse in water Fume hood, 2 times | | ON | 0 | 20 | 50 | OFF | |
| 48 | Rinse in water | 00:00:40 | ON | 250 | 20 | 50 | OFF | |
| 49 | Rinse in water | 00:00:40 | ON | 250 | 20 | 50 | OFF | |
| 50 | Osmium ON | 00:02:00 | ON | 100 | 20 | 50 | AUTO | 1% OsO4 in aqueous solution |
| 51 | Osmium OFF | 00:02:00 | OFF | 0 | 20 | 50 | AUTO | |
| 52 | Osmium ON | 00:02:00 | OFF | 100 | 20 | 50 | AUTO | |
| 53 | Osmium OFF | 00:02:00 | OFF | 0 | 20 | 50 | AUTO | |
| 54 | Osmium ON | 00:02:00 | OFF | 100 | 20 | 50 | AUTO | |
| 55 | Osmium OFF | 00:02:00 | OFF | 0 | 20 | 50 | AUTO | |
| 56 | Osmium ON | 00:02:00 | OFF | 100 | 20 | 50 | AUTO | |
| 57 | Rinse in water Fume hood, 2 times | | ON | 0 | 20 | 50 | OFF | |
| 58 | Rinse in water | 00:00:40 | ON | 250 | 20 | 50 | OFF | |
| 59 | Rinse in water | 00:00:40 | ON | 250 | 30 | 50 | OFF | |
| 60 | Uranyl acetate ON | 00:02:00 | ON | 100 | 40 | 50 | AUTO | 1 % uranyl acetate in aqueous solution, filtered |
| 61 | Uranyl acetate OFF | 00:02:00 | OFF | 0 | 40 | 50 | AUTO | |
| 62 | Uranyl acetate ON | 00:02:00 | OFF | 100 | 40 | 50 | AUTO | |
| 63 | Uranyl acetate OFF | 00:02:00 | OFF | 0 | 40 | 50 | AUTO | |
| 64 | Uranyl acetate ON | 00:02:00 | OFF | 100 | 40 | 50 | AUTO | |
| 65 | Uranyl acetate OFF | 00:02:00 | OFF | 0 | 40 | 50 | AUTO | |
| 66 | Uranyl acetate ON | 00:02:00 | OFF | 100 | 40 | 50 | AUTO | |
| 67 | Rinse in water Fume | | ON | 0 | 40 | 50 | OFF | |
| 68 | Rinse in water | 00:00:40 | ON | 250 | 40 | 50 | OFF | |
| 69 | Rinse in water | 00:00:40 | ON | 250 | 40 | 50 | OFF | |
| 70 | Lead aspartate ON | 00:02:00 | ON | 100 | 50 | 60 | AUTO | 0.02 M lead nitrate and 0.03 M aspartic acid (pH 5.5), filtered |
| 71 | Lead aspartate OFF | 00:02:00 | OFF | 0 | 50 | 60 | AUTO | |
| 72 | Lead aspartate ON | 00:02:00 | OFF | 100 | 50 | 60 | AUTO | |
| 73 | Lead aspartate OFF | 00:02:00 | OFF | 0 | 50 | 60 | AUTO | |
| 74 | Lead aspartate ON | 00:02:00 | OFF | 100 | 50 | 60 | AUTO | |
| 75 | Lead aspartate OFF | 00:02:00 | OFF | 0 | 50 | 60 | AUTO | |

| | | | | | | | |
|---|---|---|---|---|---|---|---|
| 76 | Lead aspartate ON | 00:02:00 | OFF | 100 | 50 | 60 | AUTO |
| 77 | Rinse in water Fume | | ON | 0 | 20 | 50 | OFF |
| 78 | Rinse in water | 00:00:40 | ON | 250 | 20 | 50 | OFF |
| 79 | Rinse in water | 00:00:40 | ON | 250 | 20 | 50 | OFF |
| 80 | 20% Ethanol | 00:00:40 | ON | 250 | 20 | 50 | OFF |
| 81 | 20% Ethanol | 00:00:40 | ON | 250 | 20 | 50 | OFF |
| 82 | 50% Ethanol | 00:00:40 | ON | 250 | 20 | 50 | OFF |

**Supplementary Table 2. All gene counts from the zebrafish forebrain choroid plexus. Related to Figure 2.**
Each column indicates Ensemble ID, Gene name and TPM.

**Supplementary Video 1. Labeling both the blood plasma and the brain ventricles by intravenous injection of fluorescent dyes. Related to Figure 4.**
The 3D reconstructed video of the *Gt(fabp7b:2A-Gal4vp16);Tg(uas:NTR-mCherry);Tg(uas:starmaker-GFP)* line injected 10 kDa Alexa-647 fluorescent dye at 17 dpf. The confocal stack used in Supplementary Figure 4 was utilized for 3D reconstruction.
